# Reinforcement learning as an intermediate phenotype in psychosis? Deficits sensitive to illness stage but not associated with polygenic risk of schizophrenia in the general population

**DOI:** 10.1101/668939

**Authors:** M Montagnese, F Knolle, J Haarsma, JD Griffin, A Richards, P Vertes, B Kiddle, PC Fletcher, PB Jones, MJ Owen, P Fonagy, ET Bullmore, R Dolan, NSPN Consortium, M Moutoussis, I Goodyer, GK Murray

## Abstract

**Background:** Schizophrenia is a complex disorder in which the causal relations between risk genes and observed clinical symptoms are not well understood and the explanatory gap is too wide to be clarified without considering an intermediary level. Thus, we aimed to test the hypothesis of a pathway from molecular polygenic influence to clinical presentation occurring via deficits in reinforcement learning.

**Methods:** We administered a reinforcement learning task (Go/NoGo) that measures reinforcement learning and the effect of Pavlovian bias on decision making. We modelled the behavioural data with a hierarchical Bayesian approach (hBayesDM) to decompose task performance into its underlying learning mechanisms. Study 1 included controls (*n*= 29, F|M=0.81), At Risk Mental State for psychosis (ARMS, *n*= 23, F|M=0.35) and FEP (First-episode psychosis, *n*= 26, F|M=0.18). Study 2 included healthy adolescents (*n*= 735, F|M= 1.06), 390 of whom had their polygenic risk scores for schizophrenia (PRSs) calculated.

**Results:** Patients with FEP showed significant impairments in overriding Pavlovian conflict, a lower learning rate and a lower sensitivity to both reward and punishment. Less widespread deficits were observed in ARMS. PRSs did not significantly predict performance on the task in the general population, which only partially correlated with measures of psychopathology.

**Conclusions:** Reinforcement learning deficits are observed in first episode psychosis and, to some extent, in those at clinical risk for psychosis, and were not predicted by molecular genetic risk for schizophrenia in healthy individuals. The study does not support the role of reinforcement learning as an intermediate phenotype in psychosis.

## 1.1 Introduction

Cognitive deficits are commonly observed in schizophrenia, including deficits in decision making and in reinforcement learning (RL, trial and error based learning from feedback). RL is a cognitive domain of interest, not only because impairments in this domain may have a direct impact on educational and occupational outcomes, but also because RL deficits may mechanistically contribute to the pathogenesis of positive and/or negative symptoms of schizophrenia and other psychoses (Frank, 2008; Deserno et al., 2013; Murray et al., 2016). RL has been suggested as a candidate process for an intermediate phenotype in schizophrenia, lying on the casual path between identified risk factors and the full clinical expression of the phenotype of illness (Kasanova et al., 2018)

Despite the strong role for genetics in the aetiology of schizophrenia (Tsuang, 2000), there is only indirect evidence that RL deficits in schizophrenia are at least partly genetic in origin. Recent evidence indicates shared genetic overlap between the genes underpinning general intellectual function and schizophrenia liability (Toulopoulou et al., 2018), and RL correlates significantly with IQ (Chen, 2015). However, much less is known concerning the genetic basis of specific cognitive deficits in schizophrenia. There is evidence that some aspects of reward processing, which is abnormal at different stages of psychosis (Murray et al., 2008; Ermakova et al., 2018), may be an intermediate phenotype in schizophrenia. For example, relatives of people with schizophrenia show altered brain activation during reward anticipation during fMRI scans (Grimm et al., 2014). Furthermore, molecular genetic risk for schizophrenia is associated with reward related brain activation: in the IMAGEN study of about 2000 14-year-olds, Lancaster et al. (2016) found that schizophrenia polygenic risk scores were associated with striatal activation during reward anticipation. If this altered brain activation is manifest in the altered ability to learn about rewards and reward-related decision making, then we expect that reward-based reinforcement learning behaviour should also be related to polygenic risk for schizophrenia.

If RL is an intermediate phenotype for schizophrenia, individuals who are at increased clinical risk of developing psychosis (At Risk Mental States ARMS), might show a deficit in RL, but of lesser severity than individuals with the full illness phenotype. It is not yet established whether schizotypal traits or clinical risk for psychosis are associated with altered RL. Recent evidence has suggested that patients at clinical risk for psychosis show subtle subcortical prediction error abnormalities during RL (Ermakova et al., 2018), but whether these neural deficits are associated with the behavioural deficits are not clear.

There is some suggestion that RL abnormalities may be particularly prominent in certain patient groups. For example, reward-related RL deficits are particularly prominent in patients with negative symptoms, possibly contributing to the pathogenesis of such symptoms (Gold et al., 2012). Further support for the link between RL deficits and negative symptoms comes from computational modelling studies that tried to tease apart the different learning mechanisms involved. Albrecht et al., (2016) administered a Go/NoGo RL task (Guitart-Masip et al., 2012) to a group of chronic schizophrenia patients. Patients showed impaired Pavlovian biases, a tendency to seek reward with action invigoration and avoid punishment with action suppression, possibly suggesting a reduction of those mechanisms in the striatal regions and a disruption in communication between these striatal areas and the prefrontal cortex. The influence of Pavlovian biases on RL has not been studied in first episode psychosis (FEP) or clinical risk for psychosis, and it is not known whether deficits in RL differ across different stages of psychotic illness or are linked to use of medication. The effects of Pavlovian biases on learning and decision making are of interest in relation to pathogenesis of psychiatric symptoms (Moutoussis et al., 2018) and in decision-making in everyday life (Hunt et al., 2016).

If RL is an intermediate phenotype in schizophrenia, we hypothesised to find RL deficits in FEP patients, in ARMS individuals, and in members of the general population with a raised molecular genetic risk for the disorder. Further, we would expect that RL performance would relate to trait schizophrenia measures in the population. We thus studied RL in a group of FEP patients, ARMS individuals, and healthy individuals. In several hundred healthy individuals, we examined whether their performance on a RL task related to their molecular genetic risk for schizophrenia and to their psychopathology. We hypothesised that impairments in RL would relate to trait level manifestations of subclinical positive and negative symptoms. We combined standard measures of learning with a computational psychiatry analysis approach (Teufel & Fletcher, 2016; Redish & Gordon, 2016), as it offers the possibility of developing rigorous and testable models of behaviour that can contribute to our understanding of how abnormal neurobiological substrates become expressed in clinical phenotypes.

## 2.1 Methods and Materials

### 2.1.1 Participants

#### Clinical study

We recruited three groups of participants aged 17 to 35 (mean age 22.8 years): *n*= 23 participants for the ARMS group, *n*= 26 FEP patients and *n*= 29 Controls. FEP participants were recruited from the Cambridge First Episode Psychosis service, CAMEO. ARMS participants belonged to the APS group and were recruited through CAMEO, through advertisements at University Counselling Services, and from existing local research databases; ARMS status was confirmed using the CAARMS interview Comprehensive Assessment of At Risk Mental States (CAARMS), as used in the EDIE-II trial (Morrison et al., 2012) – all ARMS participants met CAARMS attenuated psychotic symptoms criteria. Medication details can be found in *Table 12* in the Supplementary Section. Controls were recruited thorough advertisement in Cambridgeshire and through existing University of Cambridge research databases. Exclusion criteria: current or past history of neurological disorder or trauma, currently or recently participating in a clinical trial of an investigational medical product, learning disability, or not satisfying standard MRI safety exclusion criteria, including pregnancy. The latter requirement was due to the fact that a subset of volunteers had MRI scans, reported elsewhere (Whittaker et al., 2016). Past or current treatment for a mental health problem was an exclusion criterion for controls. The project received ethical approval from the National Research Ethics Service. Written informed consent was signed by all participants; if they were below 16 years of age, then written parental consent was also required. Further demographic information can be found in *Table 1* below.

**Table 1.**
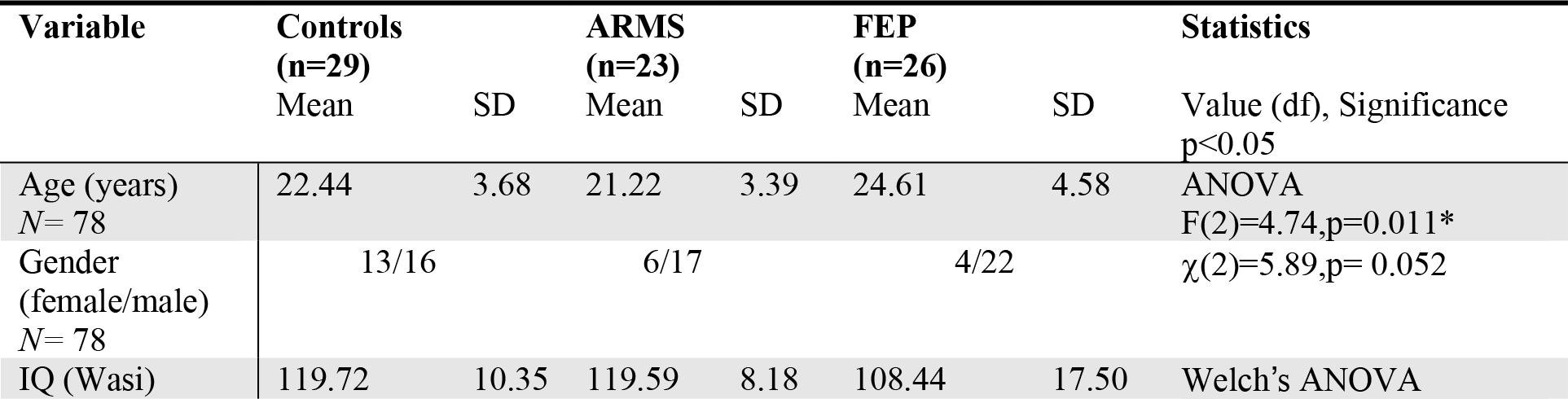
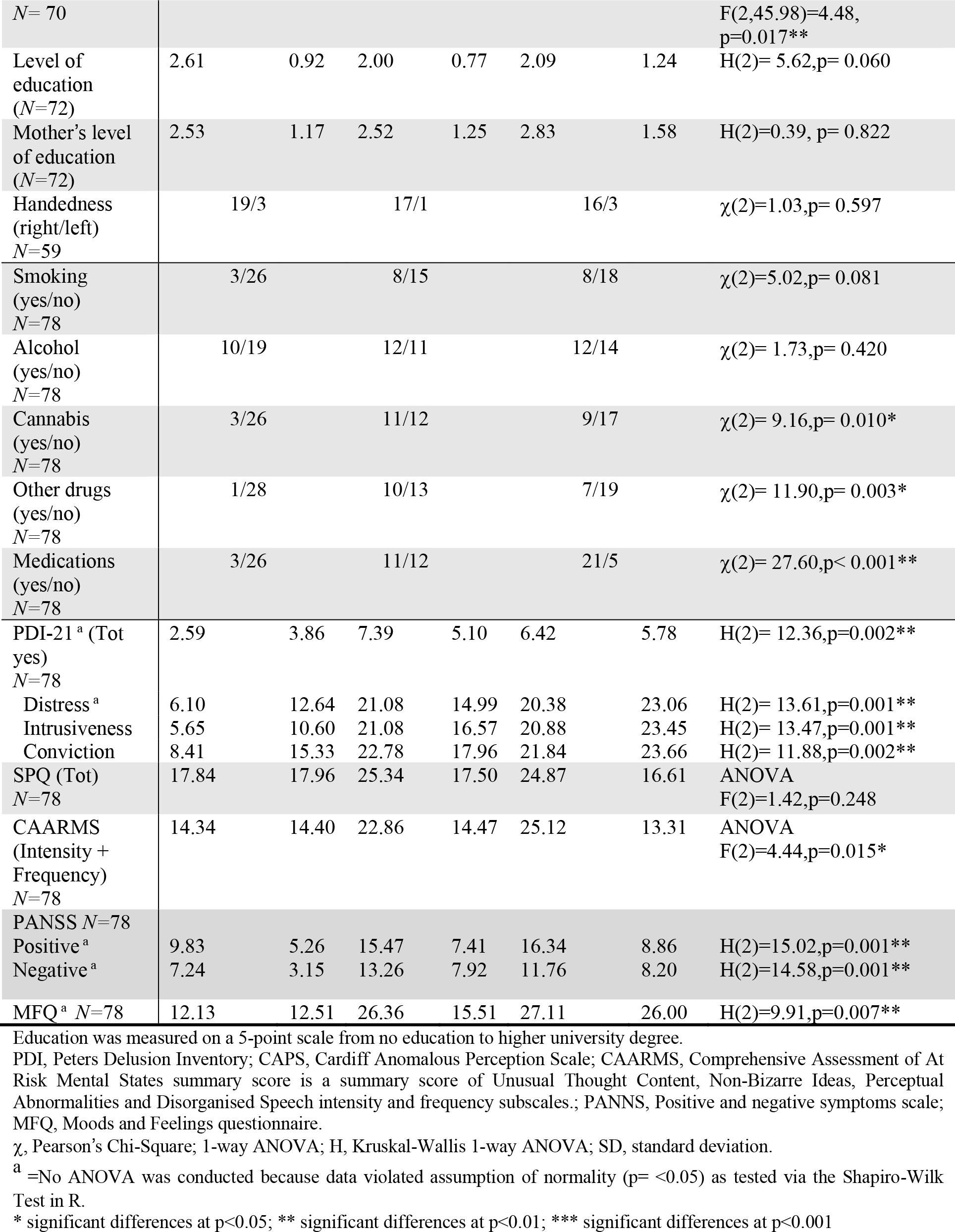
Demographic information for the Clinical Study

#### Healthy adolescent volunteer study

*N*= 785 participants took part (mean age 18.6 years, SD= 2.96; F|M=1.06) and underwent cognitive RL testing. Participants were recruited from General Medical Practice lists as a sampling frame as well as by direct advertisement so as to represent the UK population in this age range (Kiddle et al, 2017). Inclusion criteria were age 14 to 24 years old, able to understand written and spoken English, living in Greater London or Cambridgeshire & Peterborough, being willing and able to give informed consent for recruitment into the study cohort and consent to be re-contacted directly. Exclusion criteria were as described above for controls in the clinical study. A detailed analysis of reinforcement performance in these participants is available in Moutoussis et al., (2018), which does not address molecular genetics or schizotypal traits. See Table 2 below for full demographic information.

**Table 2.** Table of demographics for the Healthy Adolescent Study

| Variable | Baseline assessment<br>(n=735) |  |
| --- | --- | --- |
|  | Mean | SD |
| Age (years) | 18.60 | 2.96 |
| Gender (female/male) | 379/356 |  |
| IQ<br>(from WASI vocab and matrix combined)<br>2 missing | 111.01 | 11.32 |
| Level of education | 2.05 | 1.39 |
| Mother's level of education | 2.03 | 1.34 |
| Father's level of education | 1.77 | 1.36 |
| Handedness (100=right/0=left)<br>12 missing | 64.86 | 48.58 |
| Smoking (yes/no)<br>1 missing | 137/597 |  |
| Alcohol (yes/no)<br>8 missing | 476/251 |  |
| Cannabis (yes/no)<br>3 missing | 85/647 |  |
| Other drugs (yes/no)<br>5 missing | 43/687 |  |
| Medications (yes/no)<br>12 missing | 94/629 |  |
| MFQ (Tot) -1 missing | 16.57 | 11.62 |
| PLIKS (Tot yes) | 0.31 | 0.78 |
| SPQ (Tot) | 19.51 | 11.98 |
| SHAPS (Tot no) – N= 533 | 0.59 | 1.48 |

Education was measured on a 5-point scale from no education to higher university degree.
MFQ, Moods and Feelings Questionnaire; PLIKS (Unusual experience, Hallucination); SPQ, Schizotypal Personality Questionnaire; SHAPS, Snaith-Hamilton Pleasure scale.

### 2.1.2 Psychopathology measures

The participants in the Clinical study were administered: the Comprehensive Assessment of At Risk Mental States (CAARMS) (Yung et al., 2005), providing operational criteria for identification of clinical risk for psychosis; the Mood and Feelings Questionnaire (MFQ) subset of the Young People Questionnaire (YPQ) (Costello & Angold, 1988) to measure depressive symptoms; the Positive and Negative Symptoms Scale (PANSS)(Kay et al., 1987); to measure schizotypy they were administered the 21-items Peters Delusions Inventory (PDI-21) (Peters et al., 2004) and the Schizotypal Personality Questionnaire (SPQ); IQ was measured from combining the scores of two subscales of the Wechsler Abbreviated Scale of Intelligence (WASI), namely the Vocabulary and Matrix subtests. The healthy adolescent participants were administered the following: MFQ; PLIKS (Psychosis-Like Symptoms) to measures unusual experiences, hallucinations and delusions (Zammit et al., 2008). The Schizotypal Personality Questionnaire (SPQ)(Raine, 1991) to measure schizotypy. The SPQ was later scored according to the novel subscales provided by Davies (2017); the Snaith Hamilton Pleasure Scale (SHAPS)(Snaith et al., 1995) to measure some aspects of anhedonia (higher scores reflect higher values of anhedonia); IQ was measured from the WASI, the same way as in the Clinical study.

### 2.1.3 Reinforcement learning task

All participants were assessed on a modified version of a traditional Go/NoGo RL task, developed by Guitart-Masip et al., (2012) that provides several measures of RL (*Figure 1)*. The task involved the presentation of four fractal images 36 times each, for a total of 144 trials across the 4 conditions. The order of the stimuli was random and each cue was presented for 800ms, followed by cross-hair in the middle of the screen for 250-3500ms. Then there was a target detection task showing a circle on either side of the screen for a maximum time of 800ms, during which time the participant had to make a button press response (Go) or not (NoGo). The Go response was given via pressing a keyboard button on the side on which the cue was presented (right or left), then the probabilistic outcome was shown. Possible outcomes were: a green arrow upward for wins (£0.5), a red one downwards for losses (-£0.5) and a yellow horizontal bar for neutral outcomes (£0). For the reward conditions, only positive or neutral outcomes were possible, while for the losses conditions participants could experience either a loss or a neutral outcome. Importantly, these outcomes were probabilistic on a 80:20 schedule. Overall, there were four conditions depending on the cue presented at the start of the task: two *Pavlovian congruent* conditions requiring to press the button to get a reward (*Go-to-win*) or to not press the button to avoid losing (*NoGo-to-avoid-losing*); two *Pavlovian Incongruent* conditions requiring to either not press the button to get a reward (*NoGo-to-win*) or to press the button to avoid losing (*Go-to-avoid-losing*). (see Supplementary Material for details).

**Figure 1.**
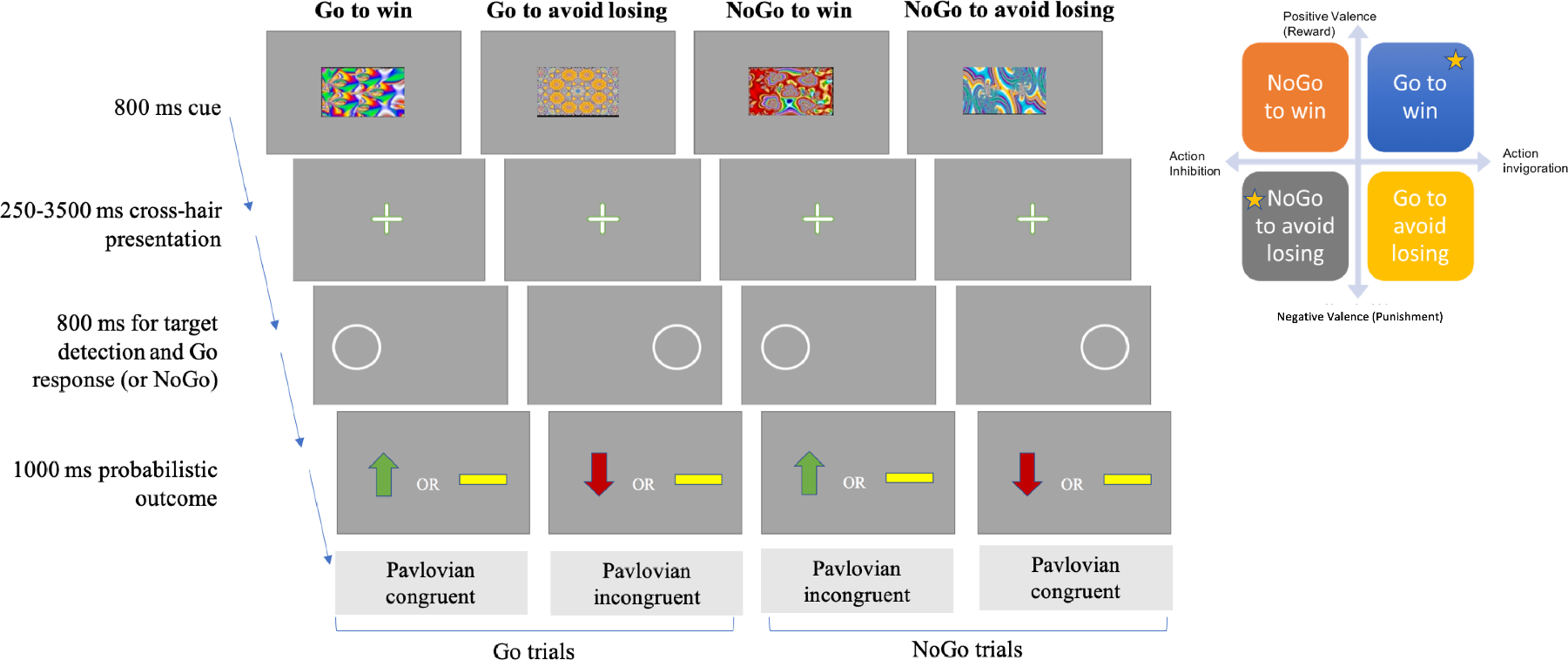
Experimental paradigm schematic. Figure adapted from *Guitart-Masip et al., (2012) and Moutoussis et al 2018.* Top-right figure shows a graphical representation of the four conditions of the modified Go/NoGo task crossing valence (y-axis) and action (x-axis). Yellow stars mark the Pavlovian congruent conditions, while the other two are the Pavlovian incongruent ones.

### 2.1.4 Computational modelling: hBayesDM

Behavioural performance on the Go/NoGo task was calculated by summing scores for the task conditions, and by modelling latent task variables using the hBayesDM package (hierarchical Bayesian modeling of Decision Making tasks) for R (version 0.5.0 on MacOS High Sierra version 10.13.1) developed by Ahn et al. (2017). We used this approach to generate posterior distributions of the parameters characterising task performance to improve the balance of within-subject and between-subject random effects, whilst also taking into account within-subject variability and group-level similarities (O’Callaghan et al., 2017). Full information on the details of the modelling parameters and model fitting and comparison can be found in the Supplementary Material. “Model 4” was the best model (lowest LOOIC) for both cohorts of participants and included the following parameters: lapse rate (random errors), learning rate, Go bias (tendency to make a response), Pavlovian bias (tendency to make a response to stimuli associated with reward and withhold a response to stimuli associated with punishment), sensitivity to reward, sensitivity to punishment. HBayesDM produces posterior distributions of modelled parameters for each individual; for each individual we selected the median of the distribution to take forward into statistical analysis to compare groups or examine within group correlations.

### 2.1.5 Polygenic risk score calculation

Participants in the healthy adolescent study participants were drawn from a larger sample of over 2000 adolescents on whom genetic data were acquired from by saliva sample (Kiddle et al., 2017). Genotyping was carried out by the Cambridge Bioresource on an Affymetrix chip array, yielding genotype at 507,968 SNPs for subjects. Quality control and imputation was performed. The parameters for retaining SNPs were: SNP missingness < 0.01 (before sample removal); SNP Hardy-Weinberg equilibrium (P > 10^−6^) and minor allele frequency MAF > 0.01. Final statistical analyses were carried out on *n=*390 participants of European ancestry for whom both adequate genotype and RL data were available. See *Figure 4* in Supplementary Material for a detailed flowchart of excluded participants. The generation of the PRS was based on the methods described by the *International Schizophrenia Consortium* (2009). Polygenic scores were calculated for each individual using the PLINK (version 1.9) score command. Scores were created by adding up the number of risk alleles for each SNP, i.e. single nucleotide polymorphism, which took the value of 0,1, or 2 and weighted by the logarithm of its odds ratio for schizophrenia from the results reported in Pardinas et al., (2018): the meta-analysis of the CLOZ-UK sample and the Psychiatric Genomics Consortium PGC2 schizophrenia dataset (Jones et al., 2016). The scores used were generated from a list of SNPs with a GWAS training-set P<.05 threshold, as this is the threshold that has been suggested to capture maximal schizophrenia liability (Schizophrenia Working Group of the Psychiatric Genomics Consortium 2014; Pardiñas et al. 2018).

### 2.1.6. Statistical analyses

In the clinical study, group differences on behavioural task performance were examined by one-way analysis of variance (ANOVAs). For the group differences in modelled parameters we run both an ANOVA with the median values, as well as an ANCOVA with the median values and subject-level uncertainty for model fit as a covariate (See Supplementary Material). Sensitivity analyses were run to compare the results of these group comparisons with and without outliers (defined as being outside of 1.5*Interquartile Range). Spearman Rank Order correlation coefficients were used investigate the relationships between task performance and clinical measures at each group level. Despite the group differences in IQ in the clinical study, since matching for education and IQ could yield a non-representative sample of patients, and given that both the participants’ own level of education and their maternal levels of education were not significantly different from controls, we did not match ARMS and FEP for IQ and, like Albrecht et al., (2016), we did not use IQ as a covariate for the statistical analyses carried out.

In the healthy adolescent study, the relationships between task performance (behavioural and modelled) and clinical measures were examined by Spearman Rank Order correlation coefficients (*n*= 735). Standard multiple regression analysis was first used to test whether PRS at P-threshold 0.05 predicted learning rate as measured by the computational model (chosen as the main outcome variable given the robust evidence in the literature showing learning deficits in patients with schizophrenia). Covariates included age, sex and the first five primary component analysis factors for ancestry. *N*= 5 participants were excluded as outliers (see Supplementary Material), with a final sample of *n*= 390. To test if the PRS scores predicted the other aspects of task performance, standard multiple regression analyses were then run for each of the other cognitive variables of interest. False Discovery Rate (Benjamini-Hochberg) correction was applied to control for the expected proportion of falsely rejected hypotheses and to gain power (Benjamini & Hochberg, 1995). Further, Bayesian linear regressions were also performed in JASP to compare the likelihood of the task performance data under models with, versus without, schizophrenia polygenic risk score.

## 3.1 Results

### 3.1.1 Clinical study

All groups showed the classic pattern of better performance in the Pavlovian congruent conditions. A one-way ANOVA across the three groups revealed significant differences in overall performance across groups (F(2, 75)= 4.61, p= .013). In terms of overall performance (percent for best outcome) on the four GNG conditions, all groups showed better performance in the Pavlovian congruent conditions compared to the Pavlovian incongruent ones. FEP performed significantly worse than controls and ARMS in the Punishment conditions (Go-to-avoid losing and NoGo-to-avoid-losing) and also worse than controls on the easier Go-to-win condition. See *Figure 2* **below** and the descriptive statistics in *Table 4* in the Supplementary Material. To further explore the possible effect of antipsychotic medication on our results, we then subdivided the FEP group into two different sub-groups: one of individuals who did not take antipsychotics (FEP-*n=* 11) and one with those taking antipsychotics, (FEP+ *n=* 15). The overall group difference in performance for the FEP group was particularly prominent in the FEP+ subgroup. (See Supplementary Material).

**Figure 2.**
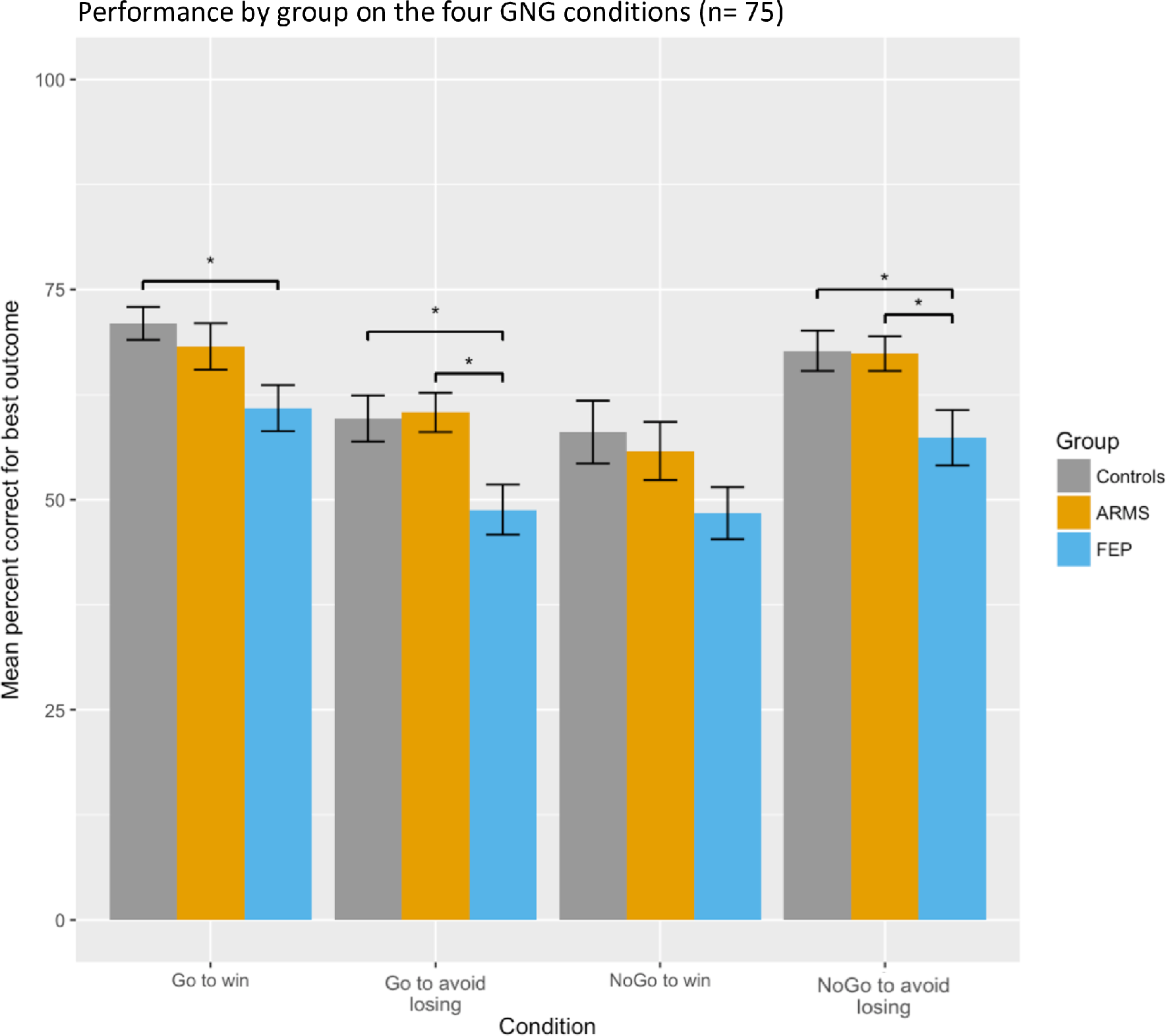
Group differences in overall performance (percent for best outcome) on the four GNG (Go/NoGo) conditions. Controls n=29, ARMS (At-Risk for Mental Health) n=23 and FEP (First-episode psychosis) n=26. Error bars indicate standard error of the mean. Stars indicate significant t-test group differences at p<0.05 after ANOVA testing.

We found group differences for all of the six modelled parameters (latent variables of task performance; Figure 3). Results were essentially unchanged after accounting for subject-level uncertainty in model fitting with the ANCOVA. See Supplementary Material for the ANCOVA analysis and details of the post-hoc comparisons for all analyses.

**Figure 3.**
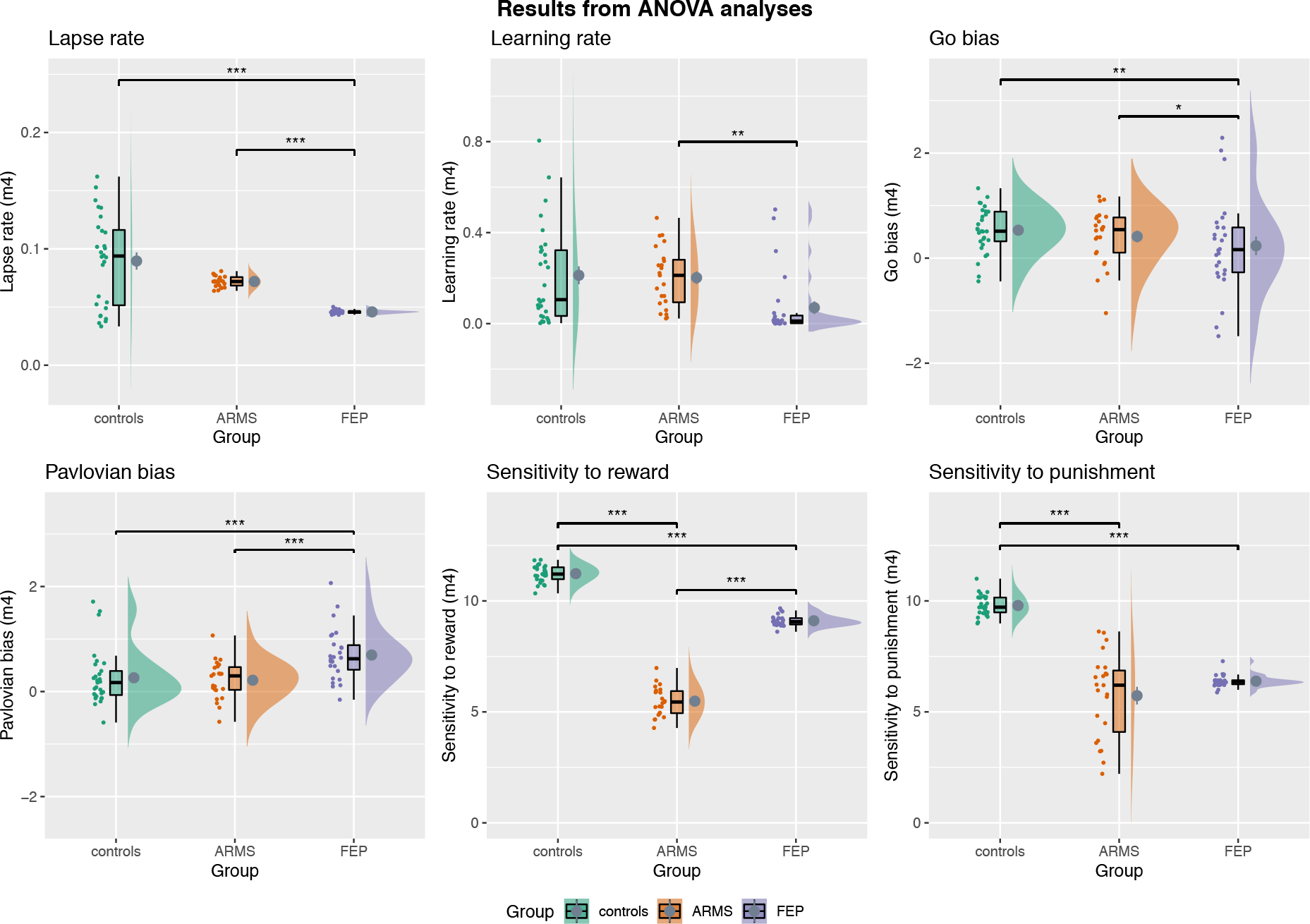
ANOVA analysis of group differences of modelled parameters. Controls, ARMS (At-Risk for Mental Health) and FEP (First-episode psychosis). For Go Bias and Pavlovian Bias, values >0 indicate the presence of such bias, those <0 indicate the opposite. Horizontal back bar=Median; mean=grey circle. Whiskers indicate the interquartile range and the cloud plot shows the probability distribution of the data. Significant results from the Bonferroni or Games-Howell corrected post-hoc analyses are shown after outliers removal (*p<.05, **p<.01, ***<.001).

Results from the Spearman correlational analyses investigating possible relationships between task performance and clinical measures for each group can be found in *Figure 7* in the Supplementary Material. Overall, for ARMS the negative symptoms (measured with the PANSS) positively correlated with learning rate. The SPQ and some of its subscales were negatively correlated with reward sensitivity and positively to punishment sensitivity. For the FEP group, learning rate negatively correlated with the positive symptoms (measured via the PANSS) and Pavlovian bias negatively correlated with both total SPQ and with its subscale of social anhedonia.

### 3.1.2 Healthy adolescent study

The pattern of performance in the healthy adolescent study is reported in detail in Moutoussis et al., (2018). In brief, there were, as expected, significant differences in performance across conditions, with better performance on the Pavlovian congruent conditions compared to the Pavlovian incongruent ones, and similar patterns for the learning curves. The Spearman correlational analyses on the Healthy Adolescent group showed a moderate negative correlation between the modelled parameters of Pavlovian bias and that of learning rate. Moutoussis et al., (2018) reported that there were no significant associations between task indices and mood. Our behavioural results (*Figure 8* in Supplementary Material) indicate weak positive correlations between the Go bias parameter and SPQ tot (r=.13, p=.01), as well as with two SPQ subscales tapping on social anxiety and eccentricity (r=.13, p=.01 and r=0.10 p=.04). The SPQ subscale reflecting anomalous experiences and beliefs was weakly negatively correlated with the sensitivity to reward in the task (r=-.14, p< .001). The sensitivity to punishment was weakly negatively associated with the SPQ subscale of paranoid ideation (r=-.15, p<.001) and with the PLIKS (r=-.11, p=.03).

The results from the standard multiple regression analysis between PRS at P-threshold 0.05 and the learning rate parameter (with age, sex, first five primary component analysis factors for ancestry as covariates) was not statistically significant: R^2^=.005, F(8, 381)=.218, p=.988, adjusted R^2^=-.016, Unstandardized B Coefficient=.001 (Standard error=.008, t-value=.065, p=.988). Standardized Beta coefficient (β)=.003. See *Figure 9* in the Supplementary Material. Results for the other main cognitive variables of interest are summarised in *Table 3* below in ascending order of adjusted significance p-value. Overall, after corrections, no significant results were found.

**Table 3.** Summary of the results from the standard multiple regressions carried out with PRS P-threshold of 0.05 before and after False discovery rate (FDR) correction. P-values are shown in order of ascending order of adjusted significance. (*p<.05, **p<.01, ***<.001). SE=Standard error of the unstandardized coefficient.

| Cognitive variable of interest (IV) | Unstandardized regression coefficient and SE | Standardized coefficient ($\beta$) | t-value | Significance of regression coefficient for the PRS Score and the cognitive variable of interest | Significance after False discovery rate correction |
| --- | --- | --- | --- | --- | --- |
| NoGo-to-avoid-losing % | -1.027(.652) | -.082 | -1.575 | .116 | .528 |
| Sensitivity to punishment (median) | -.540(.364) | -.077 | -1.483 | .150 | .528 |
| Lapse rate (median) | .001(.003) | .017 | .321 | .176 | .528 |
| Pavlovian bias (median) | -.002(.023) | -.004 | -.067 | .381 | .820 |
| Go bias (median) | -.005(.031) | -.008 | -.152 | .672 | .820 |
| Sensitivity to reward (median) | .579(.402) | .076 | 1.442 | .719 | .820 |
| NoGo-to-win % | .768(1.044) | .038 | .735 | .463 | .820 |
| Go-to-avoid-losing % | -.251(.723) | -.018 | -.347 | .729 | .820 |
| Go-to-win % | .118(.559) | .011 | .211 | .833 | .833 |

We also run Bayesian linear regression analyses, comparing a model with PRS to a null model including age, gender and the first five PCA components of ancestry as covariates. Results can be found in *Table 13* in the Supplementary Material. The null model with the covariates out-predicted the model that contained the main predictor of interest for all task-related variables and thus the results converge with what found in the standard multiple regression analyses.

### 4.1 Discussion and Conclusions

In the Clinical study, all groups showed better performance in the Pavlovian congruent conditions compared to the Pavlovian incongruent ones. We found group differences in behavioural and modelled performance on the task, with FEP performing worse than the other two groups. Further to this, and contrary to what was expected, FEP performed relatively better on the Pavlovian congruent conditions compared to the Pavlovian incongruent ones.

There were significant group differences in all modelled parameters. FEP had generally higher Pavlovian bias than both ARMS and Controls. The sensitivity to reward differed across all groups, with ARMS having the lowest one, followed by FEP and then by Controls. The sensitivity to punishment differed across the groups, being lower for ARMS compared to controls, and also significantly reduced in FEP compared to controls; this is at odds with prior findings in chronic schizophrenia patients (Gold et al., 2008), and adds to evidence that reinforcement learning deficits appear to differ in early psychosis compared to chronic schizophrenia.(Chang et al., (2016)), The finding of a higher Pavlovian bias in first episode psychosis patients compared to controls is in contrast with the findings from Albrecht et al., (2016) in chronic illness, who are older and have more negative symptoms. This might be attributable to the progression of the disease which, alongside an extensive use of antipsychotics (Scherer et al., 2004), may be linked to the worsening of deficits in RL - the neural substrate of which is thought to involve the striatum. In turn, this might have the effect of weakening the Pavlovian biases, which is also linked to striatal dysregulation, and result in the pattern observed in Albrecht’s study. Further support for this possibility can be seen in our supplementary material, as the FEP+ on antipsychotic medication have a relatively lower Pavlovian bias compared to those not taking antipsychotic medications (FEP-). The ARMS group are more similar to controls than to FEP in lapse rate, learning rate, Pavlovian and go biases, but differ from controls in sensitivity to reward and punishment. If confirmed in larger samples, this finding may indicate that sensitivity to reward and punishment are the aspects of RL to be first affected in the earliest stages of psychotic illness. We recently showed evidence of mild midbrain abnormalities in prediction error signalling in ARMS, but intact cortical function (Ermkavoa et al 2018). Other tasks might reveal learning deficits in ARMS that could be detected by the current task. For further investigations longitudinal follow ups of ARMS and FEP patients, and randomised placebo controlled trials, are necessary.

In the Healthy Adolescent study, the pattern of overall performance on the RL task is the same as that of controls from the Clinical study, thus showing that, in the general population, individuals learn the Pavlovian congruent conditions more easily and have more difficulties with the incongruent ones. When correlating task performance and clinical measures of psychopathology, we found some evidence of weak associations between task performance and schizotypy. In the Healthy Adolescent study, the sensitivity to reward was negatively correlated with the schizotypy subscale tapping on anomalous experiences and beliefs, and the sensitivity to punishment was negatively correlated with the PLIKS, which also measures unusual experiences. For ARMS, higher negative symptoms were linked to better learning rate but the sensitivity to reward was worse as the schizotypy level increased for these patients, overall suggesting a link between schizotypy and impaired reward-learning. Worse punishment sensitivity was only linked to a decrease in the SPQ subscale of disorganised speech.

A similar trend linking clinical symptoms and impaired reward-related learning was found in the FEP group, where the more severe the positive symptoms, the worse was the learning rate and the more impaired was the performance on the reward related Go-to-win condition. Interestingly, higher schizotypy was correlated with decreased Pavlovian bias. Taken together, these results might suggest a link between impaired reward-related learning and schizotypy in clinical psychosis and in the healthy population.

Our results show that PRSs for schizophrenia in the general population do not predict performance on this specific RL task. There are multiple possible explanations for this, which cannot be disentangled in the current study. The first possible explanation is that the PRS for schizophrenia does not specifically bear on the cognitive domain of RL, which could be more associated with illness itself rather than illness risk. The second explanation is that the regression analyses were underpowered to detect any small polygenic risk effect sizes present in this sample and/or the GNG task might not have captured sufficient individual variability in performance (see Supplementary Materials for power calculation). We did not record fMRI responses during RL, which were shown to associate with schizophrenia PRS in a recent study (Lancaster et al., 2019).

For all cognitive outcomes measures, Bayesian analysis indicated the data was more likely under a model without schizophrenia polygenic risk score than one including it. Finally, the sample in the Healthy Adolescent study consisted of individuals who were partly recruited on the basis of their good health and it is possible that this lack in mental health variance might have reduced our ability to detect relationships between task performance and other traits.

In the Clinical study the groups differed significantly on age, and this could possibly be problematic when looking at group differences, as some studies point at age-related effects on RL performance (Samanez-Larkin & Knutson, 2015; Radulescu, Daniel and Niv, 2016); however, the group differences in age were only marginal, and the behavioural performance differences remained intact when controlling for this variable. The partly preserved RL performance in ARMS might simply be due to inadequate power, and larger studies will be required to examine this definitively. We acknowledge the possible influence of severe traumatic stress experiences, which was linked to increased Pavlovian biases in a previous study (Ousdal et al., 2018).

Overall, the current work makes some important contributions to the field of RL in schizophrenia. Firstly, we show that there are specific RL deficits in psychotic illness and that such deficits are sensitive to illness stage, being present in frank psychosis and to some extent in At Risk Mental States. Secondly, we show that there is an association between the RL domain of reward-related learning and psychopathology in the general population. Lastly, we found no large effects of molecular polygenic risk for schizophrenia in RL. Although power calculations indicating that a bigger sample would be required for definitive results (See Supplementary Material), the current findings do not support RL as an intermediate phenotype for schizophrenia.

## Acknowledgements & Funding

The authors are grateful to the volunteers who took part in these studies, as well as all the members of staff involved in the recruitment process. This work was supported by the Neuroscience in Psychiatry Network (NSPN) Consortium, a strategic award from the Wellcome Trust to the University of Cambridge and University College London (095844/Z/11/Z); by the Cambridge NIHR Biomedical Research Centre.

## Author roles

MMontagnese: conceptualization, methodology, formal analysis, writing (original draft preparation, review and editing); FK: conceptualization, methodology, formal analysis, supervision, writing (original draft preparation, review and editing); JH, JG: conceptualization, investigation, methodology, writing (review and editing); AR, PV, BK,: methodology, resources; writing; PCF conceptualization, methodology, writing (review and editing); PBJ, PF, ETB, RD, IG: conceptualization, project administration, funding acquisition, supervision, writing (review and editing); MO: resources; writing; MMoutoussis: conceptualization, methodology, supervision, writing (review and editing); NSPN Consortium: conceptualization, methodology, project administration. GKM, conceptualization, project administration, methodology, supervision, writing (original draft preparation, review and editing).

## Ethical standards

The authors assert that all procedures contributing to this work comply with the ethical standards of the relevant national and institutional committees on human experimentation and with the Helsinki Declaration of 1975, as revised in 2008.

