## Supplementary Material_v2 for "Reinforcement learning as an intermediate phenotype in psychosis? Deficits sensitive to illness stage but not associated with polygenic risk of schizophrenia in the general population"

_1_Department of Psychiatry, University of Cambridge, United Kingdom, _2_Wellcome Trust MRC Institute of Metabolic Science, Cambridge Biomedical Campus, United Kingdom, _3_Behavioural and Clinical Neuroscience Institute, University of Cambridge, United Kingdom, _4_ Max Planck University College London Centre for Computational Psychiatry and Ageing Research, United Kingdom, _5_Wellcome Centre for Human Neuroimaging, University College London United Kingdom, _6_Cambridgeshire and Peterborough National Health Service Foundation Trust, Cambridge,United Kingdom, _7_Research Department of Clinical, Educational and Health Psychology, University College London, United Kingdom, _8_MRC Centre for Neuropsychiatric Genetics and Genomics, Cardiff University, UK

**Go/NoGo task**

The traditional version of this task requires participants to press a button in order to get a monetary reward (action-invigoration condition) and to withhold this motor response in order to avoid a punishment, i.e. a monetary loss (the action-inhibition condition). In the present study, we used a modified version devised by Guitart-Masip et al., (2012), whereby action (Go and NoGo) and valence (reward and punishment) are crossed in order to have four conditions. The task involved the presentation of four fractal images 36 times each, for a total of 144 trials across the 4 conditions (unlike in the (Guitart-Masip *et al.*, 2012) study where the task included 60 trials per conditions for a total of 240 trials). The order of the stimuli was random. The timeline for each condition was as follows: each cue was presented for 800ms, followed by cross-hair in the middle of the screen for 250-3500ms. Then there was a target detection task showing a circle on either side of the screen for a maximum time of 800ms, during which time the participant had to make a motor response (Go) or not (NoGo). The Go response was given via pressing a keyboard button on the side on which the cue was presented (right or left). Then, the probabilistic outcome was shown. The outcome consisted of one of 3 possible symbols: a green arrow upward for wins (£0.5), a red one downwards for losses (-£0.5) and a yellow horizontal bar for neutral outcomes (£0). For the reward conditions, only positive or neutral outcomes were possible, while for the losses conditions participants could experience either a loss or a neutral outcome. Importantly, these outcomes were probabilistic on a 80:20 schedule, meaning that in win trials only 80% of the correct motor choices were rewarded, while 20% were neutral (no reward); conversely, in the loss trials 80% of the correct inhibition of motor choices were neutral (i.e. avoided punishment), while 20% were punished. Overall, there were four trial types depending on the cue presented at the start of the task: press the button to get a reward (*Go-to-win*), do not press the button to get a reward (*NoGo-to-win*), press the button to avoid losing (*Go-to-avoid-losing*) and do not press the button to avoid losing (*NoGo-to-avoid-losing*). Just as in Guitart-Masip et al., (2012), participants were only told that the correct choice for the initial cue could either be Go or NoGo, and they were not instructed on the action contingencies, having to learn them via trial and error, nor about the probabilistic nature of the outcomes.

**Computational modeling**

**hBayesDM package**
To model the performance on the Go/NoGo task we used a computational model provided by a user-friendly R package called *hBayesDM* (hierarchical Bayesian modeling of Decision Making tasks) version 0.5.0 on MacOS High Sierra version 10.13.1. This package was developed by Ahn, Haines and Zhang (2017).
**GNG models summary.** The hBayesDM package contains four different models for the implementation of the orthogonalized task by Guitart-Masip et al., (2012), each differing in the number of parameters included. The four different computational models overall provide the following latent modelled measures thought to underpin performance on the task:
Lapse rate **(**xi or ξ) refers to the proportions of random choices made during the task, and takes values from 0 to 1, with higher values indicating a higher proportion of random choices; Learning rate (ep or ∈): shows the efficiency of learning over the trials, takes values from 0 to 1 and higher values indicate better learning; Go bias (b) reflects the tendency to press the keyboard button (making a Go response) irrespectively of the association between the action and the outcome of the initial cue; Pavlovian bias (pi or π): “this reflects a tendency to make responses that are Pavlovian congruent: that is to promote or inhibit *go* if the expected value of the stimulus is positive (appetitive) or negative (aversive), respectively” Ahn, Haines and Zhang (2017). It takes values from minus infinity to plus infinity; Effective size of reinforcement (rho or ρ) shows the sensitivity to both reward and punishment reinforcement combined; ranges from 0 to infinity; Effective size of reward reinforcement (rhoRew or ρ_rew_) shows the sensitivity to reward reinforcement only; ranges from 0 to infinity; Effective size of punishment reinforcement (rhoPun or ρ_pun_) shows the sensitivity to punishment reinforcement only; ranges from 0 to infinity.

**Model fitting procedure.** The following is the procedure required for model fitting, i.e. the determination of the free parameters that enhance the probability of obtaining the data given each specific model used (Palminteri, Wyart and Koechlin, 2017). The trial-by-trial data of the orthogonalized Go/NoGo task was prepared as a text file containing the following information and with the specific correct labels, as shown in brackets: subject identifier (*subjID*), cue number (*cue*) referring to the 4 different fractal cues presented and taking the value of 1= Go-to-win, 2= Go-to-avoid-losing, 3= NoGo-to-win, 4= NoGo-to-avoid-losing, whether the keyboard button was pressed or not (*keyPressed*, pressed=1, not pressed=0), outcome on each trial (*outcome*) that could be either 1 = reward, 0= neutral, -1= loss. Other information such as reaction time and trial number was present in the .txt file and was not used for modelling. Four .txt data files were created for these analyses, one for the U-Change including *n*=735 participants, and one for each of the groups in the Patient study(Controls, ARMS, FEP). Controls, ARMS and FEP were modelled as separate groups, as opposed to being one group, in order to account for any differences variance across groups and to avoid the results to be biased by the lack of accounting of sub-groups in the model. After setting up the .txt data file, each of the four GNG models were fitted with the data by using the following settings for the arguments of the model fitting command:

- niter = 2000; this refers to the number of iterations, including the warm-up.
- nwarmup = 1000; number of iterations used for warm-up only and is equivalent to a burn-in sample in Bayesian methods, specifying how many MCMC (Markov Chain Monte Carlo) samples to be discarded after the beginning of each chain.
- nchain = 4; this refers to the number of chains to be run, i.e. how many independent sampling sequences should be used to draw samples from the posterior distribution. In fact, given that the posterior distribution is generated from a sampling process, it is best to have multiple chains to maximise the chance of getting an actual representative posterior distribution (Ahn, Haines and Zhang *hBayesDM Reference Manual,* 2018).

The parameters calculated from each model included the mean for each parameter for each participant and these were used for subsequent analyses in the *Results* section.
Model convergence was also checked to ensure that the MCMC chains used did converge to stationary target distributions as this is important to ensure accurate parameter estimates and to ensure validity if one wants to use the calculated posterior distributions.
Results from the model comparison can be found in the *Results* section. After model fitting, posterior predictive checks were done to confirm the validity of the predictions. This procedure confirmed that the chosen models simulated the data in a way that reflected the original data inputted.

**Specifics of the Hierarchical Bayesian Analysis (HBA) steps for the calculation of hyperparameters in hBayesDM for the Go/NoGo task.** As explained by Ahn, Haines and Zhang, (2017): “the posterior inference for all models is performed with a Markov-Chain Monte Carlo (MCMC) sampling scheme”. This is an algorithm implemented in Stan programming language and involves the generation of random samples from a posterior distribution. In the current study the number of MCMC samples was 4,000 (1,000 per each of the 4 chains) and the purpose of this type of large sampling is that of making an accurate approximation of a posterior distribution from it. For further information, see the section “Performing hierarchical Bayesian Analysis with Stan” in Ahn, Haines and Zhang, (2017), *Stan reference manual* (Stan Development Team, 2018) and *Chapters 7* and *14* of *Bayesian Data Analysis* by Kruschke (2014).

**Exclusions Flowchart For The Polygenic Risk Score (PRS) Analyses**

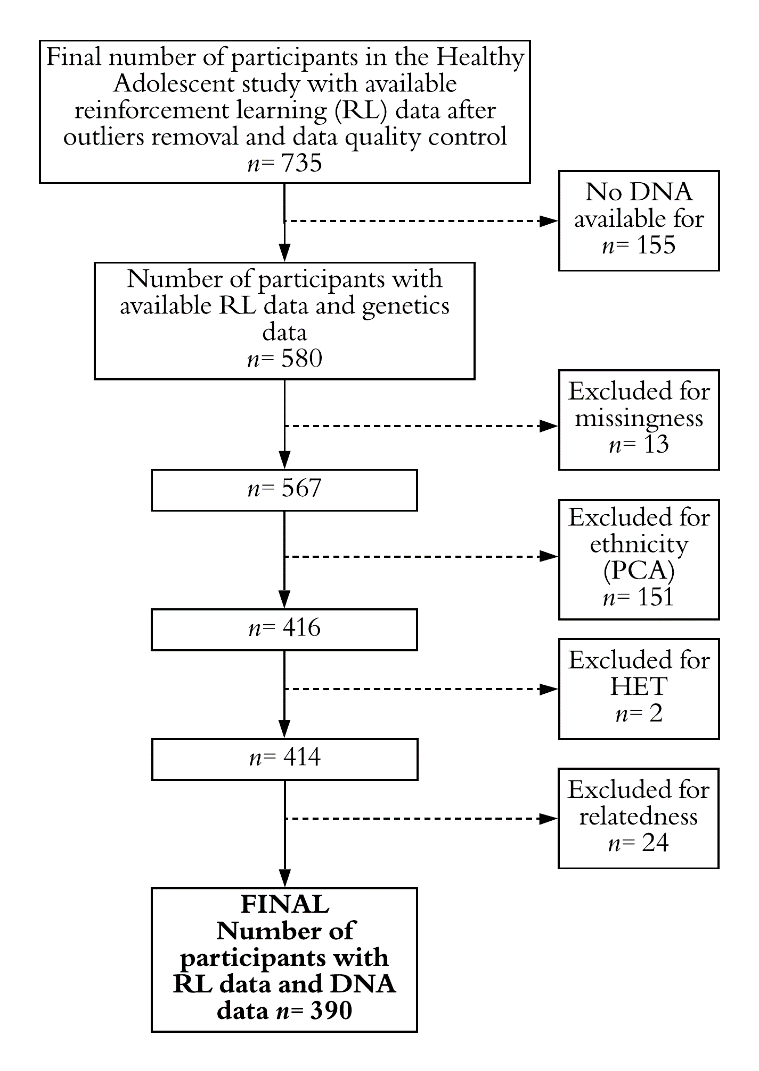

***Figure 4-*** Flowchart showing the initial number of participants who had available reinforcement learning task data (the GNG task) and DNA data, and the final number after applying different exclusion criteria. Missingness= frequency of missing data ; PCA= Principal component analysis; HET = heterozygosity; relatedness= refers to how genetically related two people are.

**Clinical Study: Statistics For Group Differences In The Behavioural Performance**

***Table 4-*** Inferential statistics of the group differences in overall performance on the four GNG conditions (percent for best outcome, *p<.05, **p<.01).

| GNG Condition | Controls  N= 29 | ARMS N= 23 | FEP N= 26 | Statistics |
| --- | --- | --- | --- | --- |
|  | Mean(SD) | Mean(SD) | Mean(SD) | ANOVA Group differences |
| Go-to-win | 70.97(10.54) | 68.23(13.26) | 60.89(13.98) | F_(2, 75)_= 4.609, **p= .013*** |
| Go-to-avoid -losing | 59.67(14.85) | 60.38(11.24) | 48.82(15.17) | F_(2, 75)_= 5.520, **p= .006**** |
| NoGo-to-win | 58.04(20.11) | 55.79(16.60) | 48.39(15.83) | F_(2, 75)_= 2.160, p= .122 |
| NoGo-to-avoid-losing | 67.72(12.91) | 67.39(9.91) | 57.37(16.81) | F_(2, 75)_= 4.873, **p= .010*** |

For the Go-to-win condition, Bonferroni post hoc analysis revealed that the mean increase in performance from FEP to ARMS (7.33, 95% CI [-1.4, 16.15]), was not statistically significant (p= .135) but the increase from FEP to Controls was (10.07, 95% CI [-1.75, 18.40], p= .012). The mean increase from ARMS to Controls was not statistically significant (2.74, 95% CI [-5.86, 11.34]), p= 1.00).

For the Go-to-avoid-losing condition, Bonferroni post hoc analysis revealed that the mean increase in performance from FEP to Controls (10.84, 95% CI [1.58, 20.11]), was statistically significant (*p*= .016), as so was the increase from FEP to ARMS (11.56, 95% CI [1.74, 21.38], *p*= .015). The mean increase from Controls to ARMS was not statistically significant (0.71, 95% CI [-8.86, 10.29]), *p*= 1.00).

Given the lack of significance for the group differences in the NoGo-to-win condition, no post-hoc analyses were run.

For the NoGo-to-avoid-losing condition, Bonferroni post hoc analysis revealed that the mean increase in performance from FEP to ARMS (10.01, 95% CI [.477, 19.56]), was statistically significant (*p*= .036) and so was the increase from FEP to Controls (10.34, 95% CI [1.34, 19.35], *p*= .019). The mean increase from ARMS to Controls was not statistically significant (0.32, 95% CI [-8.97, 9.63]), *p*= 1.00).

Further, we then looked at these group differences in behavioural performance after subdividing the FEP group into those taking antipsychotic medications (FEP+) and those not taking antipsychotics (FEP-). See *Table* *5* and *Figure* 5 below.

***Table 5-*** Inferential statistics of the group differences in overall performance on the four GNG conditions (percent for best outcome, *p<.05, **p<.01) across the four groups.

| GNG Condition | Controls  N= 29 | ARMS N= 23 | FEP- N= 11 | FEP+ N=15 | Statistics |
| --- | --- | --- | --- | --- | --- |
|  | Mean(SD) | Mean(SD) | Mean(SD) | Mean(SD) | ANOVA Group differences |
| Go-to-win | 70.97(10.54) | 68.23(13.26) | 63.38(15.35) | 59.07(13.13) | F_(3, 74)_= 3.310, **p= .025*** |
| Go-to-avoid -losing | 59.67(14.85) | 60.38(11.24) | 51.51(15.08) | 46.85(15.46) | F_(3, 74)_= 3.899, **p=.012*** |
| NoGo-to-win | 58.04(20.11) | 55.79(16.60) | 45.95(12.57) | 50.18(18.07) | F_(3, 74)_= 1.546, p= .210 |
| NoGo-to-avoid-losing | 67.72(12.91) | 67.39(9.91) | 60.10(19.65) | 55.37(14.79) | F_(3, 74)_= 3.493, **p= .020*** |

For the Go-to-win condition, Bonferroni post hoc analysis revealed that the mean increase in performance from FEP+ to Controls was (11.90, 95% CI [1.36, 22.44], p= .021).

For the Go-to-avoid-losing condition, Bonferroni post hoc analysis revealed that the mean increase in performance from FEP+ to Controls (12.82, 95% CI [.72, 24.92]), was statistically significant (p= .032), as so was the increase from FEP+ to ARMS (13.53, 95% CI [.90, 26.16], p= .029).

Given the lack of significance for the group differences in the NoGo-to-win condition, no post-hoc analyses were run.

For the NoGo-to-avoid-losing condition, Bonferroni post hoc analysis revealed that the mean increase in performance from FEP+ to Controls (12.35, 95% CI [.59, 24.10]), was statistically significant (p= .034).

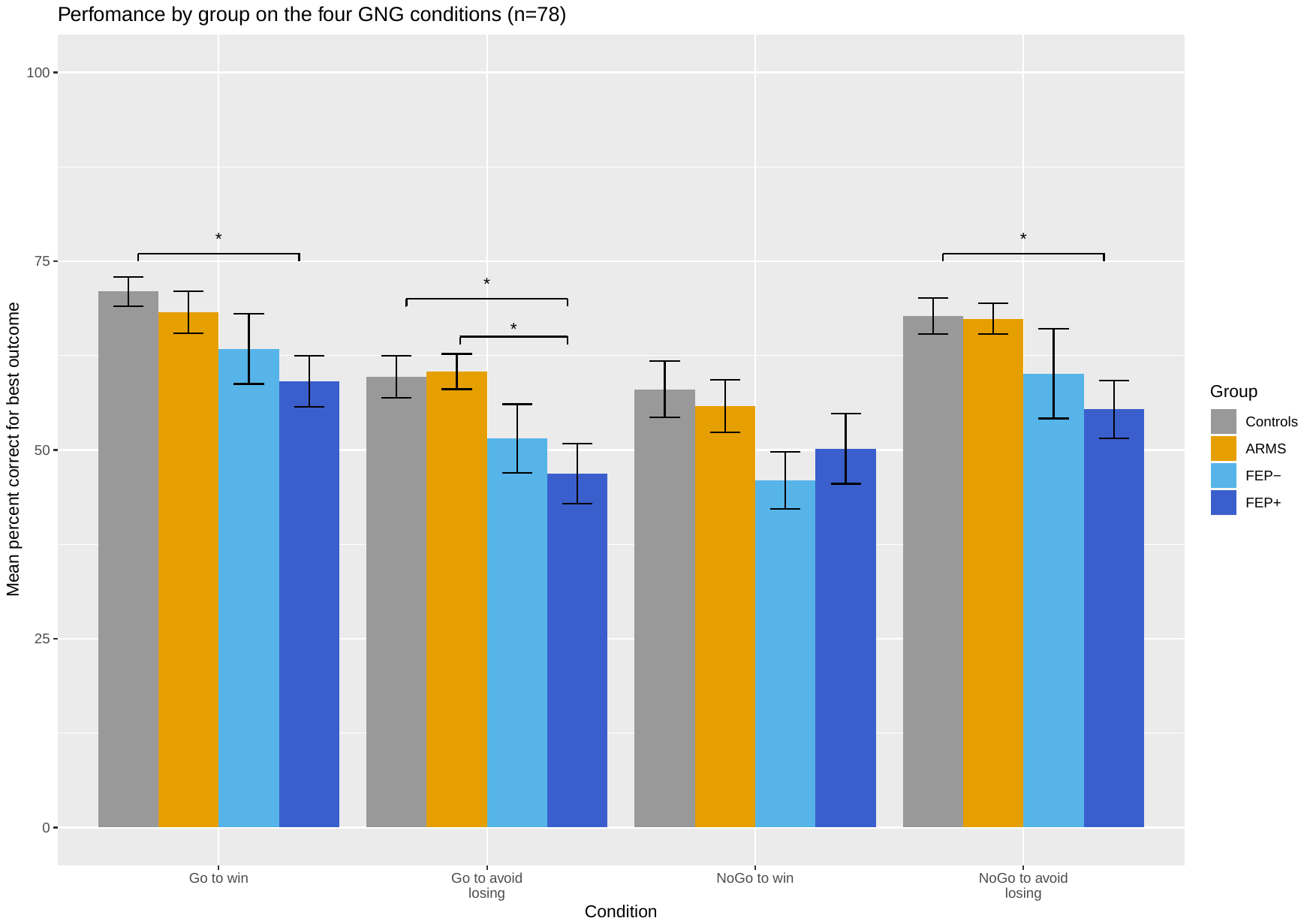

***Figure 5 -*** Group differences in overall performance (percent for best outcome) on the four GNG (Go/NoGo) conditions. Controls n=29, ARMS (At-Risk for Mental Health) n=23, FEP+ (First-episode psychosis taking antipsychotics) n=15 and FEP- (First-episode psychosis not taking antipsychotic medications) . Error bars indicate standard error of the mean. Stars indicate significant t-test group differences at p<0.05 after ANOVA testing.

**Statistics for exploratory analysis on Group*Condition Interaction**

***Table 6***- ANOVA with interaction term for Group and Condition on overall performance (percent for best outcome) (*p<.05, **p<.01, ***<.001)

| Cases | | Sum Of Squares | | Df | | | | Mean Square | | | F | | p | |
| --- | --- | --- | --- | --- | --- | --- | --- | --- | --- | --- | --- | --- | --- | --- |
| Group |  | | 6621.5 | |  | 2 |  | | 3310.77 |  | | 15.481 |  | < .001 |
| Condition |  | | 8553.0 | |  | 3 |  | | 2851.00 |  | | 13.331 |  | < .001 |
| Group ✻ Condition |  | | 173.6 | |  | 6 |  | | 28.93 |  | | 0.135 |  | 0.992 |
| Residual |  | | 64156.7 | |  | 300 |  | | 213.86 |  | |  |  |  |
| *Note.  Type III Sum of Squares* | | | | | | | | | | | | | | |

Overall, there was no significant Group*Condition interaction for performance (percent for best outcome).

**Clinical Study: Statistics For Group Differences In The Modelled Parameters**

A) **3 Groups ANOVA (after outliers removal)**

***Table 7-*** Inferential statistics for the six modelled parameters for each group (*p<.05, **p<.01, ***<.001)

ANOVA – 3 groups - after outliers removal (defined as values outside of 1.5*Interquartile Range)

| Parameters (m4) | Controls | ARMS | FEP | Statistics |
| --- | --- | --- | --- | --- |
|  | Median(IQR) | Median(IQR) | Median(IQR) | ANOVA group differences |
| Lapse rate | 0.093 (0.049-0.118)  N=28 | 0.072 (0.068-0.076)  N=23 | 0.045(0.043-0.046)  N= 22 | Welch’s F_(2, 33.940)_ = 351.407, p <.001*** |
| Learning rate | 0.099(0.028-0.305)  N = 26 | 0.212(0.089-0.286)  N=23 | 0.011(0.0006-0.502) N = 26 | F_(2, 74)_= 5.733, p= .005** |
| Go Bias | 0.512 (0.288-0.886)  N=29 | 0.552(0.111-0.7888)  N=22 | 0.120(-1.318-0.852) N = 22 | F_(2, 72)_ = 6.838, p = .002** |
| Pavlovian Bias | 0.068 (-0.099-0.297) N= 26 | 0.298 (0.026-0.477)  N=23 | 0.624(0.096-2.066) N = 24 | F _(2, 72)_ = 16.947, p <.001*** |
| Sensitivity to  reward | 11.21(10.96-11.53) N= 29 | 5.446(4.925-5.960)  N=23 | 9.064(8.612-9.667) N= 26 | Welch’s F _(2, 42.933)_ = 760.044, p<.001*** |
| Sensitivity to punishment | 9.718(9.466-10.160) N=29 | 6.203(3.703-7.002) N=23 | 6.333(5.997-6.428) N = 20 | Welch’s F_(2,35.065)_ = 750.791, p<.001*** |

**A.1) Main post-hoc analyses for the 3 groups ANOVA**
After outliers removal, there were significant group difference for all of the modelled parameters. We also did a sensitivity analysis repeating the ANOVA without outliers and the results remained essentially unchanged, with the only difference being a non-significance of the group difference for the Go bias.

For the Lapse rate parameters, Games-Howell post hoc analysis showed that the increase from FEP to ARMS was statistically significant (0.026, 95% CI [0.024, 0.029], p< .001), and so was the difference between FEP and Controls (0.044, 95% CI [0.026, 0.062], p< .001).

For the Learning rate parameter, Bonferroni post hoc analysis revealed that the increase in performance from FEP to ARMS was statistically significant ( 0.132, 95% CI [0.033, 0.230], p= .005).

For the Go bias parameter, Bonferroni post hoc analyses showed that there was a significant difference between Controls and FEP ( 0.470, 95% CI [0.139, 0.801], p= .003) and between FEP and ARMS ( 0.416, 95% CI [0.064, 0.769], p= .015).

For the Pavlovian bias, Bonferroni post hoc analyses showed that the increase from Controls to FEP was statistically significant ( 0.580, 95% CI [0.322, 0.838], p< .001), and so was the increase from ARMS to FEP ( 0.476, 95% CI [0.210, 0.742], p< .001).

For the Sensitivity to reward parameter, Games-Howell post hoc analyses showed each group differed significantly from each other. Controls differed from both ARMS ( 5.746, 95% CI [5.367, 6.125], p< .001) and from FEP ( 2.12, 95% CI [1,920, 2.326], p< .001), and FEP were also significantly different from ARMS ( 3.622, 95% CI [3.260, 3.984], p< .001).

Finally, for the Sensitivity to punishment parameter, Games-Howell post hoc analyses showed that Controls differed from both the ARMS group ( 4.062, 95% CI [3.049, 5.075], p< .001) and from FEP ( 3.484, 95% CI [3.265, 3.704], p<.001). There was no statistically significant difference between ARMS and FEP.

B) **3 groups ANOVA before outliers removal for sensitivity analyses**

***Table 8 -***Inferential statistics for the six modelled parameters for each group (*p<.05, **p<.01, ***<.001)

ANOVA – 3 groups – before outliers removal

| Parameters (m4) | Controls | ARMS | FEP | Statistics |
| --- | --- | --- | --- | --- |
|  | Median(IQR) | Median(IQR) | Median(IQR) | ANOVA group differences |
| Lapse rate | 0.093 (0.049-0.118)  N=28 | 0.072 (0.068-0.076)  N=23 | 0.045(0.043-0.046)  N= 22 | Welch’s F_(2, 35.951)_ = 328.054, p<.001*** |
| Learning rate | 0.099(0.028-0.305)  N = 26 | 0.212(0.089-0.286)  N=23 | 0.011(0.0006-0.502) N = 26 | Welch’s F_(2, 49.741)_= 7.119, p= .002*** |
| Go Bias | 0.512 (0.288-0.886)  N=29 | 0.552(0.111-0.7888)  N=22 | 0.120(-1.318-0.852) N = 22 | Welch’s F_(2,44.845)_ = 1.282, p=.287 |
| Pavlovian Bias | 0.068 (-0.099-0.297) N= 26 | 0.298 (0.026-0.477)  N=23 | 0.624(0.096-2.066) N = 24 | F _(2, 77)_ = 7.926, p=.001** |
| Sensitivity to  reward | 11.21(10.96-11.53) N= 29 | 5.446(4.925-5.960)  N=23 | 9.064(8.612-9.667) N= 26 | Welch’s F _(2, 42.933)_ = 760.044, p<.001*** |
| Sensitivity to punishment | 9.718(9.466-10.160) N=29 | 6.203(3.703-7.002) N=23 | 6.333(5.997-6.428) N = 20 | Welch’s F_(2,39.921)_ = 577.411, p<.001*** |

C) **4 Groups ANOVA after outliers removal**

***Table 9 -***Inferential statistics for the six modelled parameters for each group (*p<.05, **p<.01, ***<.001). FEP+ = First-Episode Patients taking antipsychotic medications; FEP- = First-Episode Patients not on antipsychotic medications.

ANOVA 4 groups and after outliers removal (defined as values outside of 1.5*Interquartile Range)

| Parameters (m4) | Controls | ARMS | FEP- | FEP+ | Statistics |
| --- | --- | --- | --- | --- | --- |
|  | Median(IQR) | Median(IQR) | Median(IQR) | Median(IQR) | ANOVA group differences |
| Lapse rate | 0.093 (0.049-0.118)  N=28 | 0.072 (0.068-0.076)  N=23 | 0.045(0.044-0.045)  N= 10 | 0.046(0.045-0.046) N= 12 | Welch’s F_(3, 33.681)_ = 233.991, p <.001*** |
| Learning rate | 0.099(0.028-0.305)  N = 26 | 0.212(0.089-0.286)  N=23 | 0.014(0.06-0.046) N = 11 | 0.007(0.001-0.024) N = 15 | F_(3, 74)_= 4.140, p= .009** |
| Go Bias | 0.512 (0.288-0.886)  N=29 | 0.552(0.111-0.7888)  N=22 | 0.254(-0.270-0.442) N = 8 | 0.074(-0.317-0.580) N = 14 | F_(3, 69)_ = 4.516, p = .006** |
| Pavlovian Bias | 0.068 (-0.099-0.297) N= 26 | 0.298 (0.026-0.477)  N=23 | 0.656(0.227-1.08) N = 11 | 0.579(0.486-0.785)  N = 13 | Welch’s F _(3, 73)_ = 3.977, p <.001*** |
| Sensitivity to  reward | 11.21(10.96-11.53) N= 29 | 5.446(4.925-5.960)  N=23 | 9.219(8.990-9.525) N= 11 | 8.965(8.910-9.138) N = 15 | Welch’s F _(3, 34.740)_ = 503.965, p<.001*** |
| Sensitivity to punishment | 9.718(9.466-10.160) N=29 | 6.203(3.703-7.002) N=23 | 6.248(6.196-6.345) N = 8 | 6.370(6.279-6.421) N = 12 | Welch’s F_(3,28.651)_ = 495.603, p<.001*** |

**C.1) Main post-hoc analyses for the 4 groups ANOVA**
After outliers removal, there were significant group differences for all of the modelled parameters across the 4 groups.

For the Lapse rate parameters, Games-Howell post hoc analysis showed thae following significant differences: the increase from FEP- to FEP+ (0.001, 95% CI [0.0001, 0.002], p= .022), the increase from FEP- to ARMS (0.027, 95% CI [0.024, 0.029], p< .001) and from FEP- to Controls (0.044, 95% CI [0.024, 0.064], p< .001). The increase from FEP+ to ARMS (0.025, 95% CI [0.023, 0.028], p< .001) and to Controls (0.043, 95% CI [0.023, 0.063], p< .001).

For the Learning rate parameter, Bonferroni post hoc analysis revealed a significant difference between FEP+ and ARMS (0.155, 95% CI [0.028, 0.281], p=.008).

For the Go bias parameter, Bonferroni post hoc analyses showed that there was a significant difference between Controls and FEP+ ( 0.452, 95% CI [0.027, 0.875], p= .031).

For the Pavlovian bias, Games-Howell post hoc analyses showed the following significant differences: Controls from FEP+ (.507, 95% CI [.267, .748], p< .001) and from FEP- (.666, 95% CI [.090, 1.241], p= .022), and ARMS from FEP+ (.404, 95% CI [.129, .678], p=.002)

For the Sensitivity to reward parameter, Games-Howell post hoc analyses showed each group differed significantly from each other. Controls differed from ARMS ( 5.746, 95% CI [5.328, 6.164], p< .001), from FEP- ( 2.01, 95% CI [1.723, 2.306], p< .001), and from FEP+ (2.20, 95% CI [1.975, 2.431], p< .001). ARMS differed from both FEP- (3.73, 95% CI [3.298, 4.164], p< .001) and from FEP+ (3.54, 95% CI [3.141, 3.944], p< .001).

Finally, for the Sensitivity to punishment parameter, Games-Howell post hoc analyses showed that Controls differed from ARMS ( 4.062, 95% CI [2.943, 5.181], p< .001) and from both FEP- (3.55, 95% CI [3.291, 3.808], p<.001) and FEP+ (3.44, 95% CI [3.201, 3.682], p< .001).

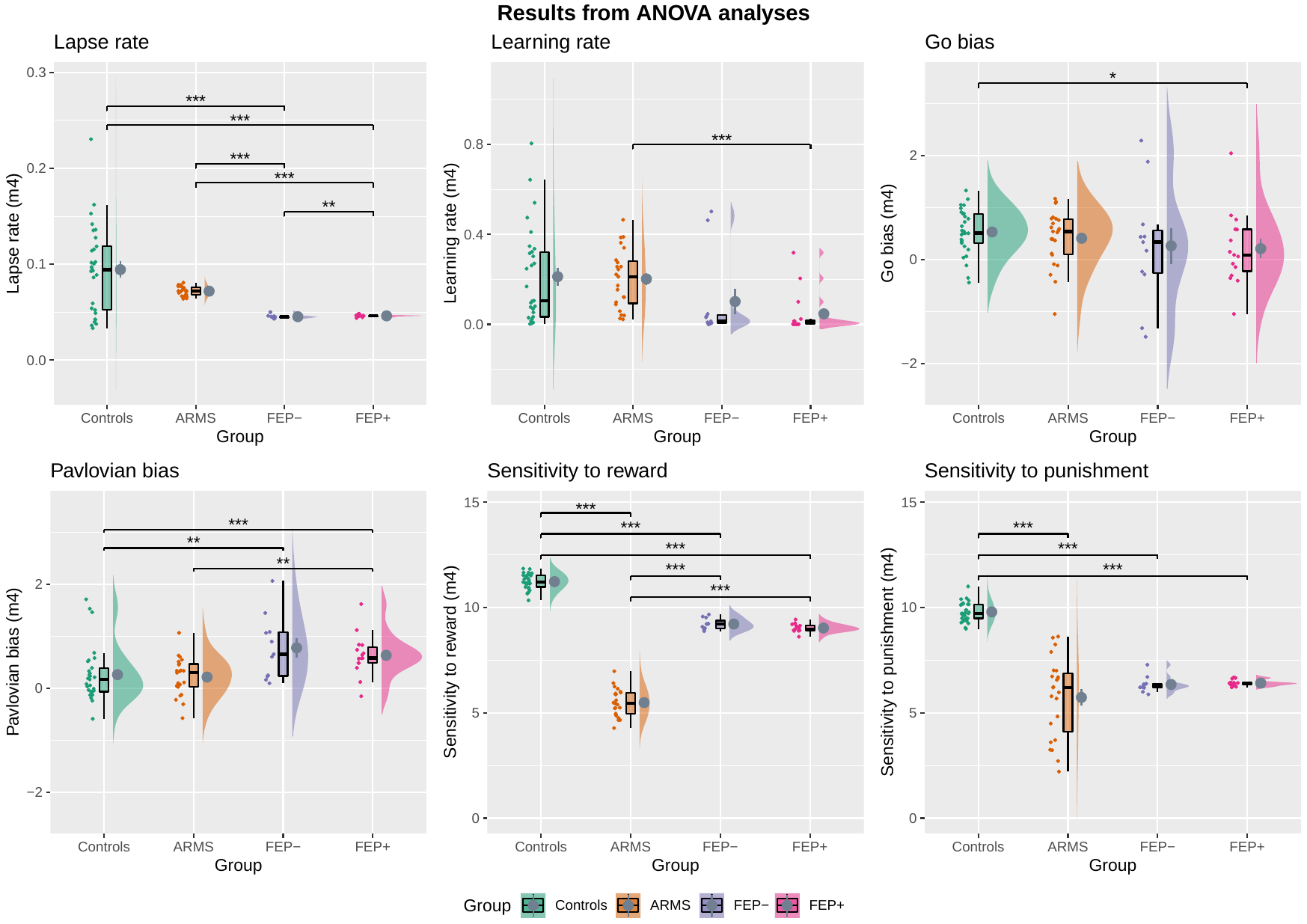

***Figure 6 –*** ANOVA analysis of group differences of modelled parameters. Controls, ARMS (At-Risk for Mental Health), FEP+ (First-Episode Patients taking antipsychotic medications), FEP- (First-Episode Patients not on antipsychotic medications). For Go Bias and Pavlovian Bias, values >0 indicate the presence of such bias, those <0 indicate the opposite. Horizontal back bar=Median; mean=grey circle. Whiskers indicate the interquartile range and the cloud plot shows the probability distribution of the data. Significant results from the Bonferroni or Games-Howell corrected post-hoc analyses are shown after outliers removal (*p<.05, **p<.01, ***<.001).

D) **3 groups ANCOVA before outliers removal for sensitivity analyses**

***Table 10 -***Inferential statistics for the six modelled parameters for each group (*p<.05, **p<.01, ***<.001)

ANCOVA 3 groups (after outliers removal)

|  | Median(IQR) | Median(IQR) | Median(IQR) | Statistics |
| --- | --- | --- | --- | --- |
|  |  |  |  | ANCOVA group differences |
| Lapse rate | 0.093 (0.049-0.118)  N=28 | 0.072 (0.068-0.076)  N=23 | 0.045(0.043-0.046)  N= 22 | F_(2,69)_ = 8.087, p<.001*** |
| Learning rate | 0.099(0.028-0.305)  N = 26 | 0.212(0.089-0.286)  N=23 | 0.011(0.0006-0.502) N = 26 | F_(2,71)_ = 1.209, p=.304 |
| Go Bias | 0.512 (0.288-0.886)  N=29 | 0.552(0.111-0.7888)  N=22 | 0.120(-1.318-0.852) N = 22 | F_(2,69)_= 4.597, p= .013** |
| Pavlovian Bias | 0.068 (-0.099-0.297) N= 26 | 0.298 (0.026-0.477)  N=23 | 0.624(0.096-2.066) N = 24 | F_(2,69)_ = .8.529, p<.001*** |
| Sensitivity to  reward | 11.21(10.96-11.53) N= 29 | 5.446(4.925-5.960)  N=23 | 9.064(8.612-9.667) N= 26 | F _(2, 74)_ =441.239, p<.001*** |
| Sensitivity to punishment | 9.718(9.466-10.160) N=29 | 6.203(3.703-7.002) N=23 | 6.333(5.997-6.428) N = 20 | F _(2, 68)_ =293.328, p<.001*** |

**D.1) Main post-hoc analyses for the 3 groups ANCOVA**

For the Lapse rate parameters, Bonferroni post hoc analysis showed that the increase from FEP to ARMS was statistically significant (0.022, 95% CI [0.08, 0.037], p= .001), and so was the difference between FEP and Controls (0.023, 95% CI [0.006, 0.039], p= .005).

For the Go bias parameter, Bonferroni post hoc analyses showed that there was a significant difference between Controls and FEP (0.408, 95% CI [0.024, 0.793], p= .040) and between FEP and ARMS (0.400, 95% CI [0.051, 0.750], p= .023).

For the Pavlovian bias, Bonferroni post hoc analyses showed that the increase from Controls to FEP was statistically significant ( 0.541, 95% CI [0.225, 0.857], p< .001), and so was the increase from ARMS to FEP ( 0.434, 95% CI [0.105, 0.764], p= .007).

For the Sensitivity to reward parameter, Bonferroni post hoc analyses showed each group differed significantly from each other. The increase from ARMS to Control was statistically significant (4.373, 95% CI [3.935, 4.811], p< .001) and so was the increase from FEP to Controls ( 2.063, 95% CI [1.856, 2.269], p< .001). The increase from ARMS to FEP was also statistically significant (2.310, 95% CI [1.884, 2.736], p< .001).

Finally, for the Sensitivity to punishment parameter, Bonferroni post hoc analyses showed that again all groups were significantly different from each other. The increase from ARMS to Controls was significant (3.789, 95% CI [3.412, 4.167], p< .001) and so was that from FEP to Controls ( 2.337, 95% CI [1.902, 2.772], p<.001). Finally, the increase from ARMS to FEP was significant as well ( 1.453, 95% CI [1.017, 1.888], p<.001)

E) **4 groups ANCOVA** after outliers removal

***Table 11 -***Inferential statistics for the six modelled parameters for each group (*p<.05, **p<.01, ***<.001)

ANCOVA 4 groups after outliers removal (defined as values outside of 1.5*Interquartile Range)

| Parameters (m4) | Controls | ARMS | FEP- | FEP+ | Statistics |
| --- | --- | --- | --- | --- | --- |
|  | Median(IQR) | Median(IQR) | Median(IQR) | Median(IQR) | ANCOVA group differences |
| Lapse rate | 0.093 (0.049-0.118)  N=28 | 0.072 (0.068-0.076)  N=23 | 0.045(0.044-0.045)  N= 10 | 0.046(0.045-0.046) N= 12 | F_(3, 68)_ = 5.330, p =002** |
| Learning rate | 0099(0.028-0.305)  N = 26 | 0.212(0.089-0.286)  N=23 | 0.014(0.06-0.046) N = 11 | 0.007(0.001-0.024) N = 15 | F_(3, 70)_= 1.216, p= .310 |
| Go Bias | 0.512 (0.288-0.886)  N=29 | 0.552(0.111-0.7888)  N=22 | 0.254(-0.270-0.442) N = 8 | 0.074(-0.317-0.580) N = 14 | F_(3, 68)_ = 3.063, p = .034* |
| Pavlovian Bias | 0.068 (-0.099-0.297) N= 26 | 0.298 (0.026-0.477)  N=23 | 0.656(0.227-1.08) N = 11 | 0.579(0.486-0.785)  N = 13 | F _(3, 68)_ = 6.132, p<.001*** |
| Sensitivity to  reward | 11.21(10.96-11.53) N= 29 | 5.446(4.925-5.960)  N=23 | 9.219(8.990-9.525) N= 11 | 8.965(8.910-9.138) N = 15 | F _(3,73)_ = 291.54, p<.001*** |
| Sensitivity to punishment | 9.718(9.466-10.160) N=29 | 6.203(3.703-7.002) N=23 | 6.248(6.196-6.345) N = 8 | 6.370(6.279-6.421) N = 12 | F_(3,67)_ = 192.9, p<.001*** |

**E.1) Main post-hoc analyses for the 4 groups ANCOVA**

For the Lapse rate parameters, Bonferroni post hoc analysis showed a significant difference between: Controls and FEP+ (0.023, 95% CI [0.001, 0.037], p= .027), ARMS and FEP- (0.022, 95% CI [0.0002, 0.041], p= .046), ARMS and FEP+ (0.023, 95% CI [0.003, 0.042], p= .013)

For the Go bias parameter, no group differences remained significant after Bonferroni correction.

For the Pavlovian bias, Bonferroni post hoc analyses showed that the increase from Controls to FEP- was statistically significant ( 0.622, 95% CI [0.110, 0.816], p< .001), and so was the increase from Controls to FEP + (0.447, 95% CI [0.102, 0.566], p= .033). There was also a significant difference between ARMS and FEP- (0.514, 95% CI [0.017, 0.703], p= .008)

For the Sensitivity to reward parameter, Bonferroni post hoc analyses showed each group differed significantly from each other, apart from no significant difference between FEP- and FEP+. Controls vs ARMS (4.383, 95% CI [1.311, 1.780], p<.001), Controls vs FEP- (2.022, 95% CI [1.738, 2.334], p<.001) and Controls vs FEP+ (2.093, 95% CI [1.596, 2.131], p<.001). The increase from ARMS to FEP- (2.361, 95% CI [0.182,0.799], p<.001) and from ARMS to FEP+ (2.289, 95% CI [0.039, 0.598], p<.001).

Finally, for the Sensitivity to punishment parameter, Bonferroni post hoc analyses showed that again all groups were significantly different from each other, with the exception of FEP- vs FEP+. The increase from ARMS to Controls was significant (3.789, 95% CI [3.297, 4.126], p< .001) and so was that from ARMS to FEP- (1.494, 95% CI [1.164, 2.384], p<.001) and to FEP+ (1.426, 95% CI [1.126, 2.184], p< .001). Controls also differed from both FEP- (2.295, 95% CI [1.344, 2.531], p< .001) and FEP+ (2.363, 95% CI [1.546, 2.566], p< .001).

**Results From The Spearman Correlational Analyses Investigating Possible Relationships Between Task Performance And Clinical Measures For Each Group in the Clinical Study**

***Figure 7****-* Correlation matrix heat-maps of behavioural performance on the four Go/NoGo task conditions, alongside modelled parameters and results from the clinical psychological measures in the Clinical study. Spearman correlation coefficient value (r) and statistical significance level of the results are shown. PANSS= Positive and Negative syndrome scale; PDI= Peter’s Delusion Index SPQ= Schizotypal Personality Questionnaire. SPQ subscales: DS= Disorganized Speech, ECC= eccentricity, SANX= Social anxiety, PI= Paranoid Ideation, SANH= Social Anhedonia, AEB= Anomalous Experiences & Beliefs.
a) Controls; b)ARMS; c) FEP

**a)**

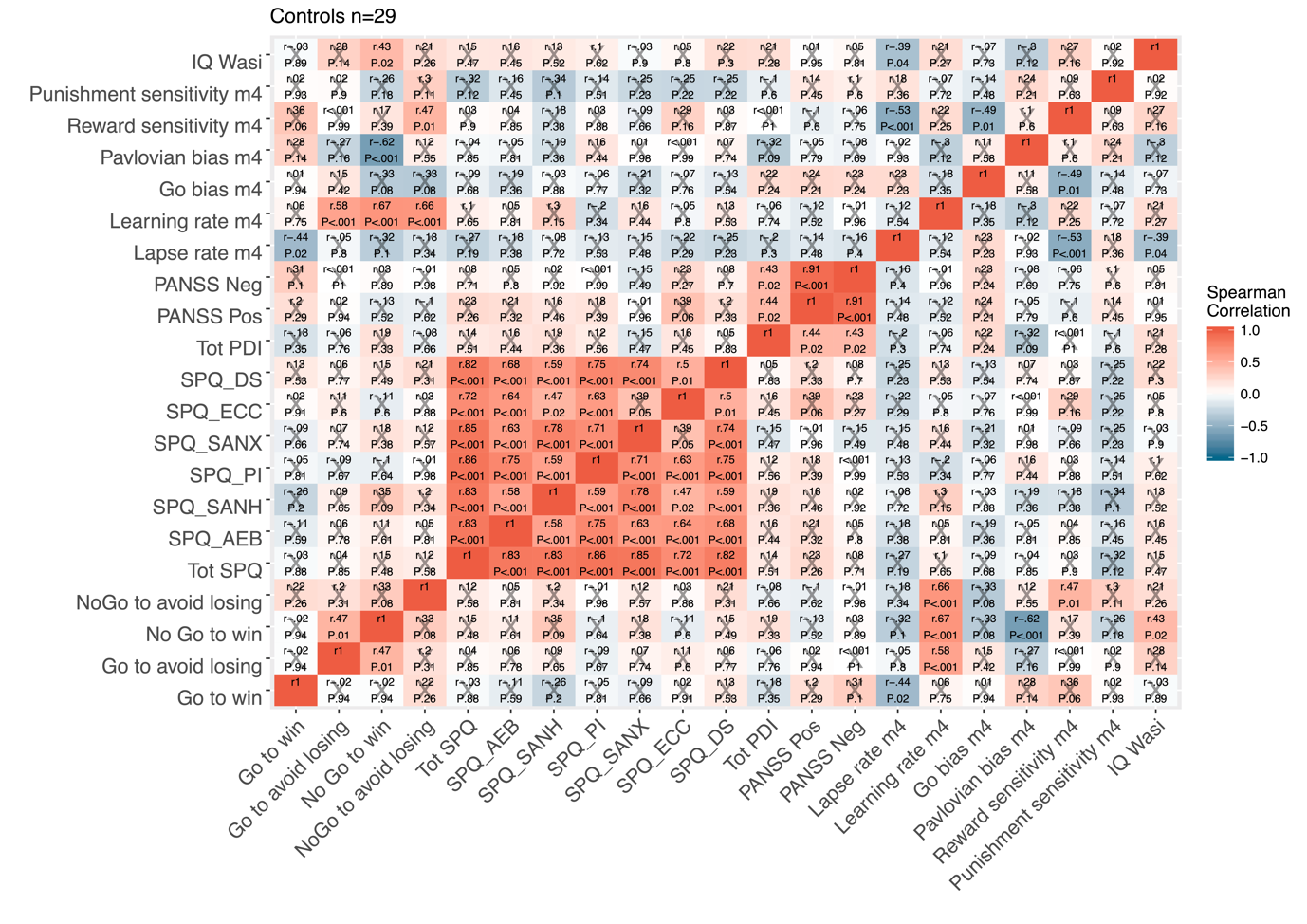

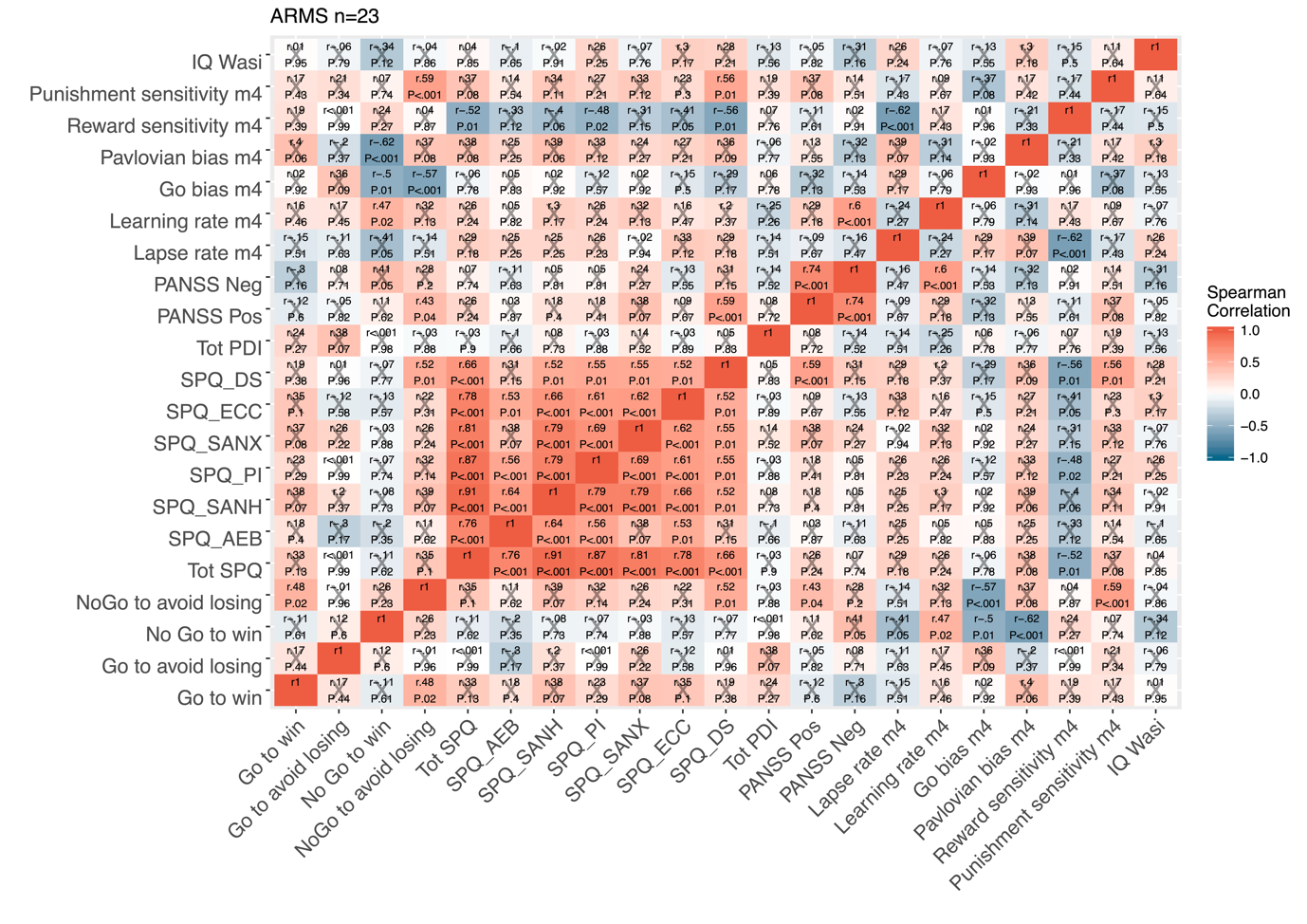

**b)**

**c)**

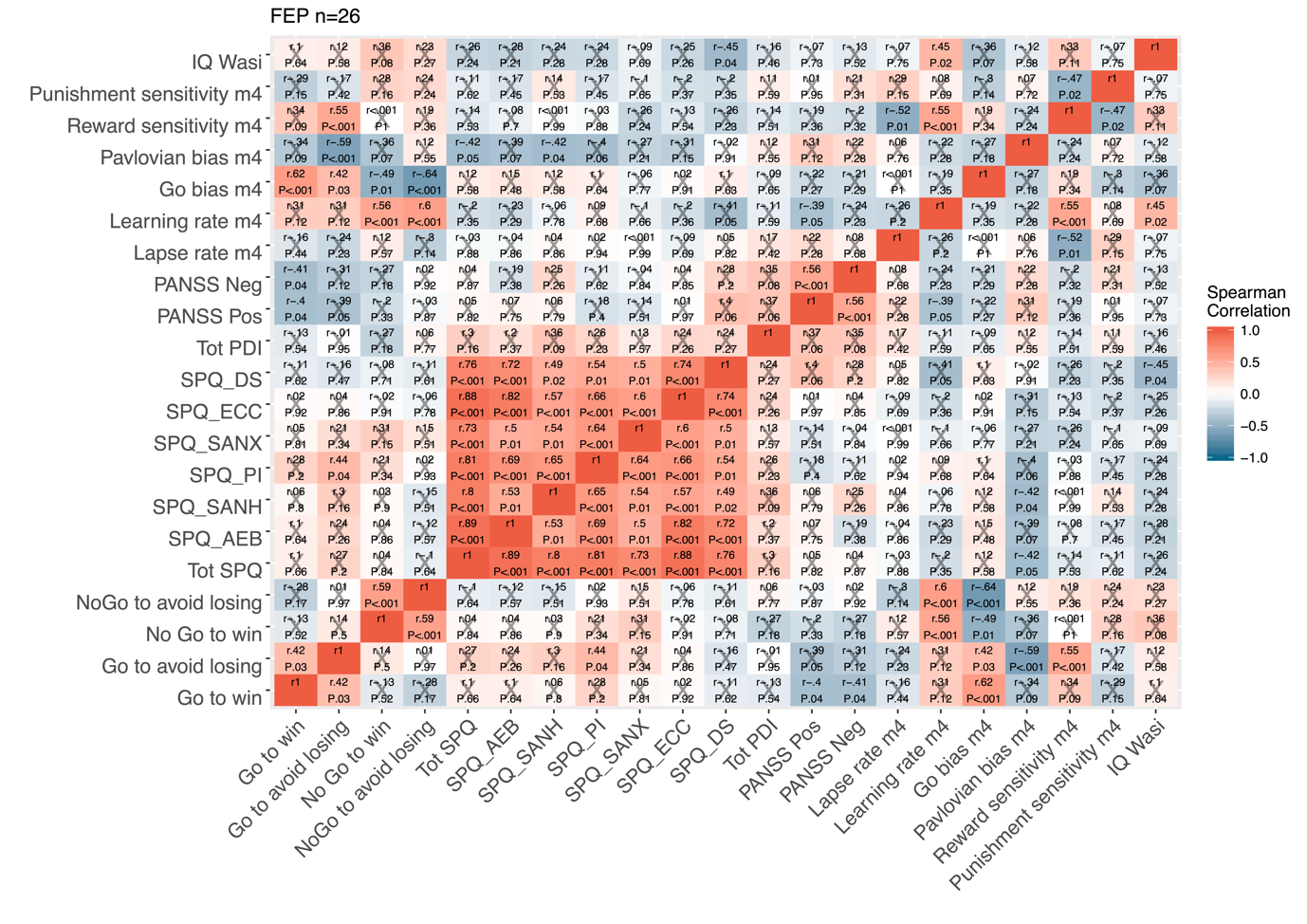

**List Of Medications For Patients In The Clinical Study**

### ***Table 12 -*** Chlorpromazine-equivalent dosages for the *Clinical study*

| Group | Antipsychotic dose | Chlorpromazine equivalent dosage |
| --- | --- | --- |
| FEP | 10 mg aripiprazole | 176.2781955 |
| FEP | 25 mg olanzapine | 692.4909639 |
| FEP | 20 mg olanzapine | 541.8885542 |
| FEP | 2 mg risperidone | 168.5689655 |
| FEP | 12,5 mg olanzapine | 315.9849398 |
| FEP | 400 mg quetiapine | 349.3347401 |
| FEP | 6 mg risperidone | 513.3965517 |
| FEP | 10 mg aripiprazole | 176.2781955 |
| FEP | 10 mg olanzapine | 240.6837349 |
| FEP | 600 mg quetiapine | 571.4582408 |
| FEP | 1 mg risperidone | 82.36206897 |
| FEP | 400 mg quetiapine | 349.3347401 |
| FEP | 15 mg aripiprazole | 364.2481203 |
| FEP | 10 mg aripiprazole | 176.2781955 |
| FEP | 3 mg risperidone | 254.7758621 |
| ARMS | 100 mg quetiapine | 16.14948912 |
| ARMS | 200 mg quetiapine | 127.2112394 |

**Results From Correlations Between Modelled And Behavioural Task Performance And The Schizotypy Measures Administered In The Healthy Adolescent Study**

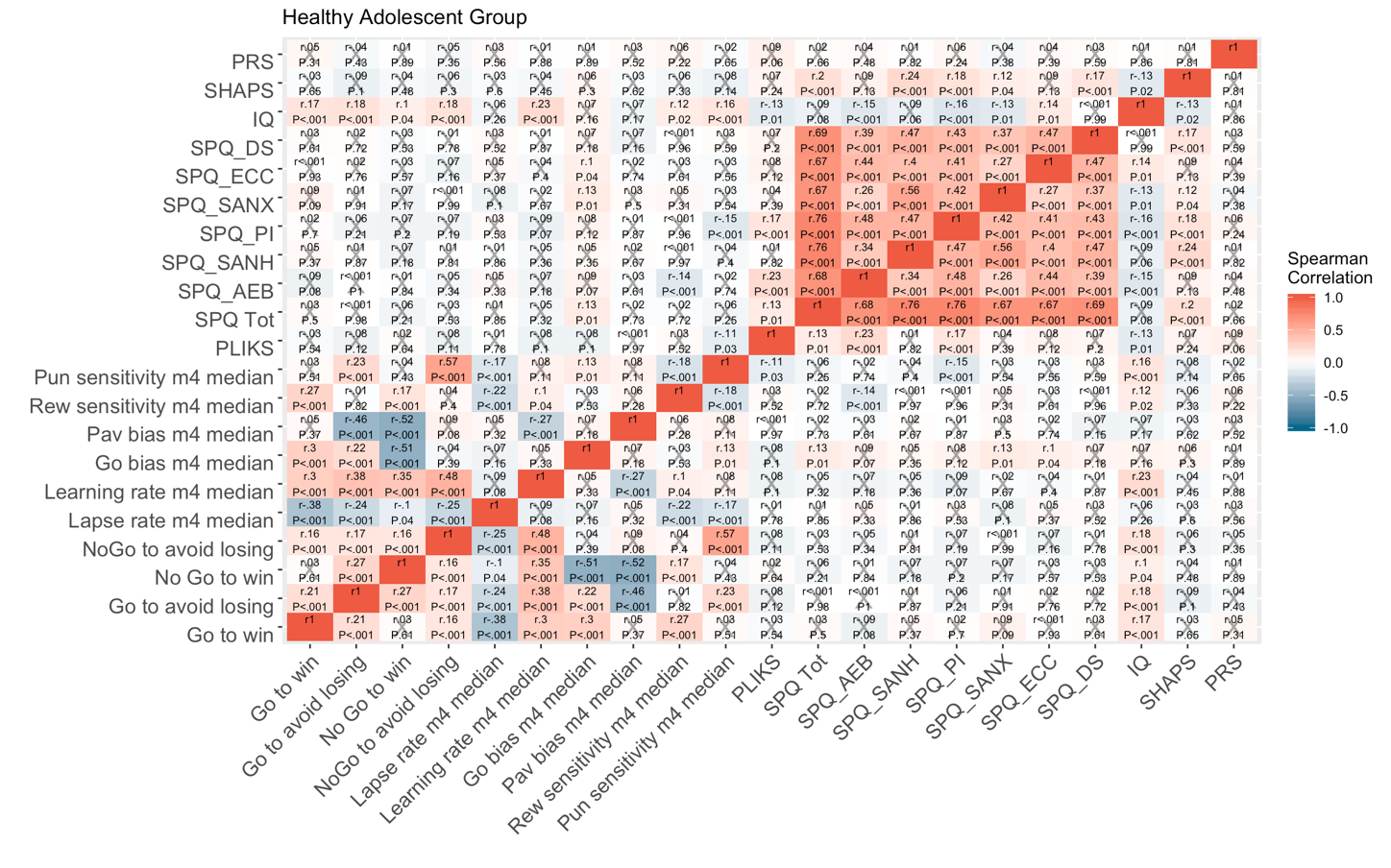

***Figure 8-*** Correlation matrix heat-maps of behavioural performance on the four Go/NoGo task conditions, alongside modelled parameters and results from the clinical psychological measures in the Healthy Adolescent study. Spearman correlation coefficient value (r) and statistical significance level of the results are shown. PANSS= Positive and Negative syndrome scale; PDI= Peter’s Delusion Index SPQ= Schizotypal Personality Questionnaire. SPQ subscales: DS= Disorganized Speech, ECC= eccentricity, SANX= Social anxiety, PI= Paranoid Ideation, SANH= Social Anhedonia, AEB= Anomalous Experiences & Beliefs.

**Multiple Standard Regression Analysis Between Modelled Parameter Of Learning Rate And PRS In The Healthy Adolescent Study.**

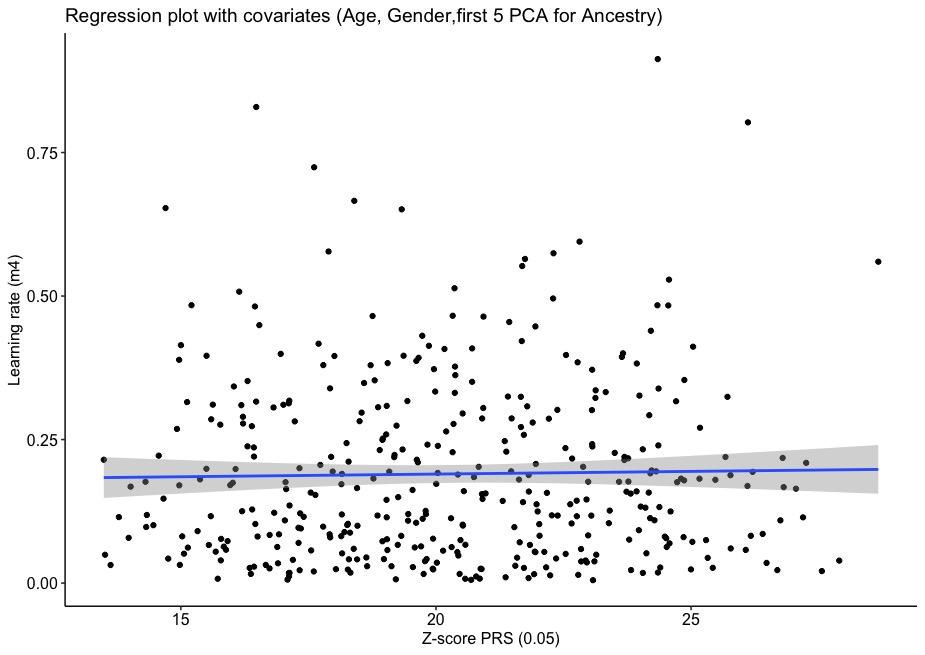

***Figure 9-*** Standard multiple regression analysis between PRS at P-threshold 0.05 and the modelled parameter of learning rate (with age, sex, first five primary component analysis factors for ancestry as covariates)

We also conducted post-hoc power calculations in order to inform future studies and examine the power of the current study to detect genetic effects on RL. Post-hoc power calculations (Soper, 2018) suggested that a sample of 390 individuals in the PRS analysis, with learning rate as the main predictor, had 0.43 power to detect an association, and therefore the analysis might have been underpowered to find any significant results. For a 0.80 power with the same observed effect size, a minimum sample of 959 individuals would have been needed to demonstrate a significant effect.

**Bayesian Regression Analyses for The Healthy Adolescent Study**

### ***Table 13*** - Bayesian Regression Analyses comparing a model with schizophrenia PRS and covariates (age, sex and the first five PCA components of ancestry) to a “null model” model 1, with the same covariates but without PRS. Implemented in JASP. BF is the Bayes Factor that shows the relative performance of Model 1 (the “null model” versus the PRS model. For each cognitive variable the first Bayes Factor is 1 (as the null model is compared to itself); the second gives the probability of the data under PRS model versus Model 1. For most cognitive variables the data is more probable under Model 1.

|  | Model Comparison |  |  |
| --- | --- | --- | --- |
| Variables | **Models** | **BF _10_** | **R²** |
| Lapse rate (m4) | Model 1 | 1.000 | 0.029 |
|  | PRS model | 0.337 | 0.029 |
| Learning rate (m4) | Model 1 | 1.000 | 0.005 |
|  | PRS model | 0.341 | 0.005 |
| Go bias (m4) | Model 1 | 1.000 | 0.015 |
|  | PRS model | 0.335 | 0.015 |
| Pavlovian Bias (m4) | Model 1 | 1.000 | 0.022 |
|  | PRS model | 0.327 | 0.022 |
| Sensitivity to reward (m4) | Model 1 | 1.000 | 0.008 |
|  | PRS model | 0.846 | 0.014 |
| Sensitivity to punishment (m4) | Model 1 | 1.000 | 0.025 |
|  | PRS model | 0.865 | 0.031 |
| Go-to-win % | Model 1 | 1.000 | 0.014 |
|  | PRS model | 0.340 | 0.014 |
| NoGo-to-win % | Model 1 | 1.000 | 0.029 |
|  | PRS model | 0.410 | 0.030 |
| Go-to-avoid-losing % | Model 1 | 1.000 | 0.004 |
|  | PRS model | 0.359 | 0.005 |
| NoGo-to-avoid-losing % | Model 1 | 1.000 | 0.023 |
|  | PRS model | 0.984 | 0.030 |

**Neuroscience in Psychiatry Network (NSPN) Consortium author list**

**Principal investigators:**

Edward T. Bullmore (CI from 01/01/2017)

Raymond J. Dolan
Ian Goodyer (CI until 01/01/2017)
Peter Fonagy

Peter B. Jones

**NSPN (funded) staff:**

Michael Moutoussis

Tobias U. Hauser

Sharon Neufeld

Rafael Romero-Garcia

Michelle St Clair

Petra Vértes

Kirstie Whitaker

Becky Inkster

Gita Prabhu

Cinly Ooi

Umar Toseeb

Barry Widmer

Junaid Bhatti

Laura Villis

Ayesha Alrumaithi

Sarah Birt

Aislinn Bowler

Kalia Cleridou

Hina Dadabhoy

Emma Davies

Ashlyn Firkins

Sian Granville

Elizabeth Harding

Alexandra Hopkins

Daniel Isaacs

Janchai King

Danae Kokorikou

Christina Maurice

Cleo McIntosh

Jessica Memarzia

Harriet Mills

Ciara O’Donnell

Sara Pantaleone

Jenny Scott

**Affiliated scientists:**

Pasco Fearon
John Suckling

Anne-Laura van Harmelen

Rogier Kievit
